# Serpina1e mediates the exercise-induced enhancement of hippocampal memory

**DOI:** 10.1101/2024.08.11.607526

**Authors:** Hyunyoung Kim, Sanghee Shin, Jeongho Han, Jong-Seo Kim, Hyungju Park

## Abstract

The exercise-induced enhancement of learning and memory is thought to be regulated by body□brain interactions via secretory proteins in the blood plasma^1,2^. Given the prominent role that skeletal muscle plays during exercise, the beneficial effects of exercise on cognitive functions appear to be mediated by muscle-derived secretory factors including myokines^3,4^. However, the specific myokines that exert beneficial effects on cognitive functions remain to be elucidated. Here, we reveal that a novel myokine, Serpina1e, acts a molecular mediator that directly supports long-term memory formation in the hippocampus. Using an *in vivo* myokine-labeling mouse model, proteomic analysis revealed that the Serpina1 family of proteins are the myokines whose levels increased the most in plasma after chronic aerobic exercise for 4 weeks. Systemic delivery of recombinant Serpina1e into sedentary mice was sufficient for reproducing the beneficial effect of exercise on hippocampus-associated cognitive functions. Moreover, plasma Serpina1e can cross the blood–cerebral spinal fluid (CSF) barrier and blood–brain barrier to reach the brain, thereby influencing hippocampal function. Indeed, an increase in the plasma level of Serpina1e promoted hippocampal neurogenesis, increased the levels of brain-derived neurotrophic factor (BDNF) and induced neurite growth. Our findings reveal that Serpina1e is a myokine that migrates to the brain and mediates exercise-induced memory enhancement by triggering neurotrophic growth signaling in the hippocampus. This discovery elucidates the molecular mechanisms underlying the beneficial effects of exercise on cognitive function and may have implications for the development of novel therapeutic interventions for alleviating cognitive disorders.

## Main

Exercise is considered a noninvasive intervention for cognitive enhancement in healthy young and aged individuals^5–7^. In addition to exercise capacity showing a strong positive correlation with academic ability in healthy humans^8^, the restorative effect of exercise on cognitive function in patients with brain disorders has been suggested^9^. This positive effect of exercise on cognitive functions is thought to be mediated by interactions between peripheral organs and the brain via the actions of secretory proteins^3,10,11^.

Skeletal muscle plays a pivotal role during exercise and is the primary organ that responds to physical activity, showing altered gene expression and protein contents^12^. Given that skeletal muscle is an active endocrine organ that releases cytokines called myokines^13^, exercise-induced myokines could be potential intermediaries of improved learning and memory through skeletal muscle–brain communication. Indeed, several *in vitro* or *ex vivo* studies identified myokines such as cathepsin B (CTSB)^3^ and Irisin^14^ as the myokines involved in exercise-induced cognitive enhancements. Despite these findings, the use of exercise interventions for cognitive and mental health issues has been limited^15^, as the detailed mechanisms of myokines involved in exercise-induced muscle-brain communication and their influence on cognitive functions are not yet fully understood. We thus aimed to identify unknown exercise-induced myokines using an organ-specific secretome-labeling tool.

### Construction of a mouse model for *in vivo* labeling of myokines

We first constructed a knock-in mouse model using an expression cassette for TurboID^16^, an efficient biotin ligase, with KDEL sequences to restrict TurboID localization into the endoplasmic reticulum lumen (ER) and a V5 tag. TurboID-V5-KDEL (TurboID-ER) expression was designed to be induced by Cre-dependent removal of the stop codon by placing a LoxP-Stop-LoxP (LSL) component between the CAG promoter and the TurboID-ER coding sequence (*LSL*-TurboID-ER; Extended Data Fig. 1). We validated the Cre-dependent expression of TurboID-ER in the knock-in mice with LSL-TurboID-ER inserted in ROSA26 by injecting Cre-expressing AAV into the *LSL*-TurboID-ER mice (Extended Data Fig. 1). The results indicated that TurboID can be expressed in Cre-expressing cells, resulting in the biotinylation of secretory proteins synthesized and secreted from Cre-expressing source cells.

To selectively label myokines released from skeletal muscles, LSL-TurboID-ER mice were crossed with *Acta1*-Cre (a skeletal muscle-specific Cre driver), resulting in ACTuR (*<u>A</u>cta1*-<u>C</u>re;*LSL*-<u>Tu</u>rboID-E<u>R</u>) mice (Extended Data Fig. 2a, b). We observed that ACTuR mice selectively expressed TurboID-ER in skeletal muscles, which enabled the selective biotinylation of myokines in ACTuR mice (Extended Data Fig. 2c, d). To detect biotinylation of skeletal muscle-derived proteins in ACTuR mice, we intraperitoneally injected biotin for 6 days and harvested several organs, such as the brain (cortex and hippocampus), liver, lung, and heart, and the hindlimb muscles. TurboID-induced biotinylated proteins were significantly distinct from endogenous biotinylated proteins (Extended Data Fig. 2c). The muscle-specific expression of TurboID-ER did not cause anxiety or depression in ACTuR mice, and no adverse effects on motor function or learning and memory were detected (Extended Data Fig. 3). Overall, we concluded that the physiological and cognitive functions of ACTuR mice were comparable to those of wild-type mice and that ACTuR mice were suitable for labeling and detecting exercise-induced myokines *in vivo*.

### Detecting aerobic exercise-induced myokines from the blood plasma of ACTuR mice

We next explored exercise-induced myokines using ACTuR mice. First, wild-type (WT) mice subjected to 4 weeks of voluntary aerobic exercise using a running wheel presented enhanced memory performance in both the contextual fear conditioning (CFC) test and the Morris water maze test (Fig. 1a, b; Extended Data Fig. 4). These enhanced memory performances were positively correlated with exercise intensity (Fig. 1c, Extended Data Fig. 4g, 4l) and lasted for three weeks (Extended Data Fig. 5b), supporting the previous notion that exercise is beneficial for learning and memory.

**Figure 1.**
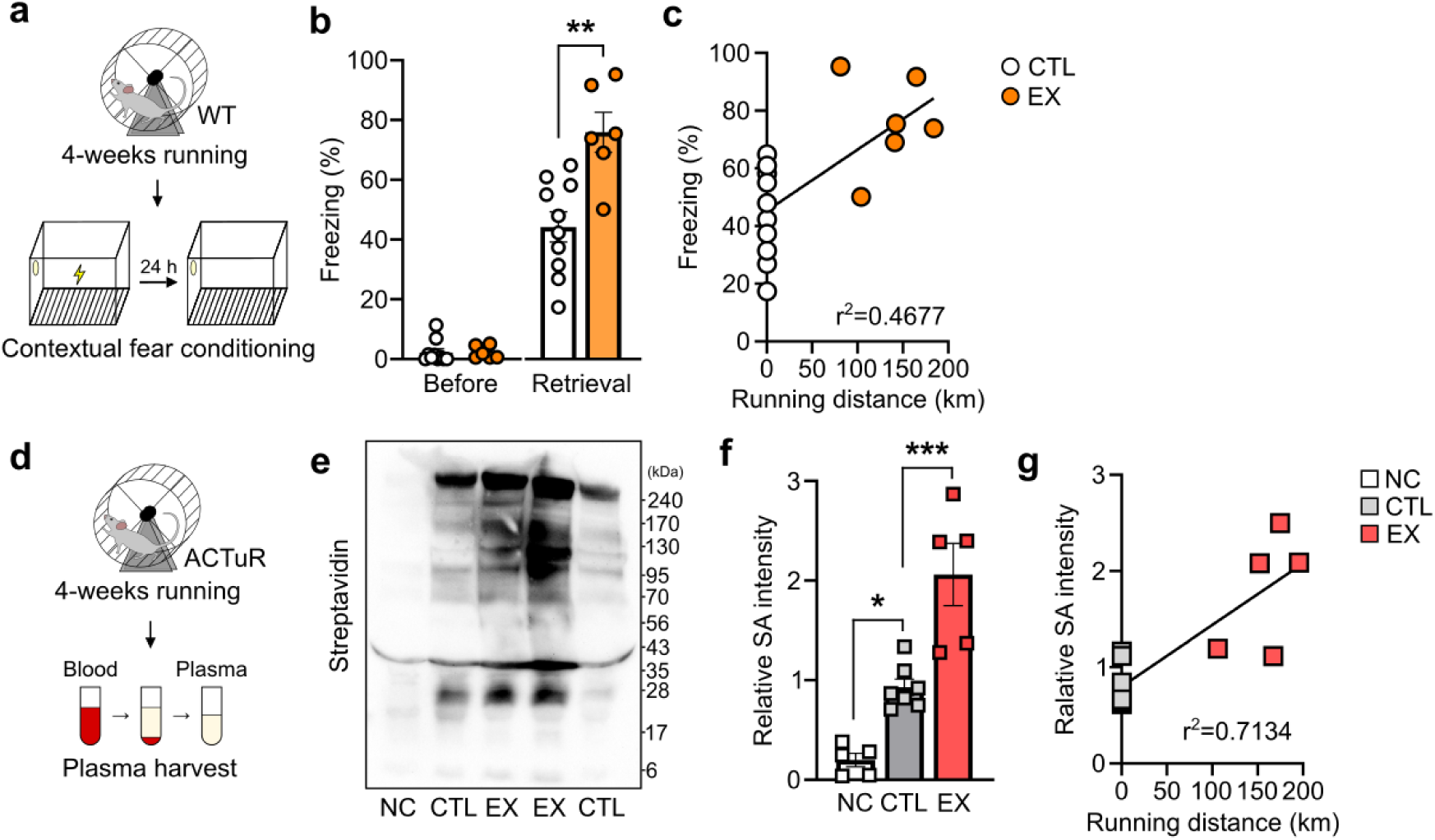
Detecting exercise-induced myokines in ACTuR mice. a. Schematic diagram showing exercise-induced alterations in hippocampal memory. b. Bar graphs depicting average percentages of freezing time (%) during the CFC test. **p=0.0019, unpaired two-tailed Student’s t test. CTL: sedentary wild-type (WT) mice; EX: exercised WT mice. c. Linear regression of the effect of running distance on freezing time. d. Schematic diagram showing the procedure for harvesting plasma from ACTuR mice after 4 weeks of exercise. e. Representative western blot results showing biotinylated proteins detected in the plasma fraction of harvested blood from ACTuR mice by staining with HRP-conjugated streptavidin (SA). NC: PBS-injected sedentary ACTuR mice; CTL: biotin-injected sedentary ACTuR mice; EX: biotin-injected exercised ACTuR mice. f. Bar graphs depicting average relative myokine levels in plasma. *p=0.0165, ***p=0.0006; one-way ANOVA with Dunnett’s post hoc test. g. Linear regression of the effect of running distance on the concentration of myokines (relative SA intensity). All the data in the bar graphs are presented as the mean ± standard error of the mean (SEM). Each dot in the graphs represent an individual tested mouse.

Using the same exercise protocol, ACTuR mice were subjected to 4 weeks of exercise to detect exercise-induced biotinylated proteins (putative myokines in ACTuR; Fig. 1d, e). As a negative control, phosphate-buffered saline (PBS), instead of biotin, was injected into ACTuR mice. Western blot analysis with horseradish peroxidase (HRP)-conjugated streptavidin revealed that the amount of plasma biotinylated proteins was increased in exercised mice compared to sedentary mice (Fig. 1f, g). Thus, these results indicate that the ACTuR mouse model is a useful tool for detecting exercise-induced plasma myokines. This exercise-induced increase in plasma myokine levels was also correlated with exercise intensity (Fig. 1h). Therefore, the beneficial effect of exercise on cognitive functions may be mediated by one or more myokines whose levels are increased by exercise.

### Identification and molecular characterization of exercise-induced myokines

To identify plasma myokines postexercise, we employed a super-resolution proximity labeling (SR-PL)^17,18^ method to identify biotinylated myokines that are secreted from skeletal muscle using liquid chromatography□tandem mass spectrometry (LC□MS/MS; Fig. 2a). Our MS/MS analysis revealed that a total of 131 proteins were biotinylated in the plasma samples, suggesting that these proteins originated from the skeletal muscles of ACTuR mice. The biotinylated proteins were quantified based on label-free quantification (LFQ) approach with good reproducibility among biological replicates (Extended Data Fig. 6a). Of the 131 proteins, we selected 78 biotinylated proteins that were found in at least 3 replicates for further analysis. These 78 biotinylated proteins overlapped well with the mouse plasma proteome (86%; Fig. 2b) and skeletal muscle proteome reference datasets (88%; Fig. 2c). Principal component analysis (PCA) showed group of clusters among replicates in each experimental group (Extended Data Fig. 6a). Therefore, we defined 78 proteins as putative myokines in plasma.

**Figure 2.**
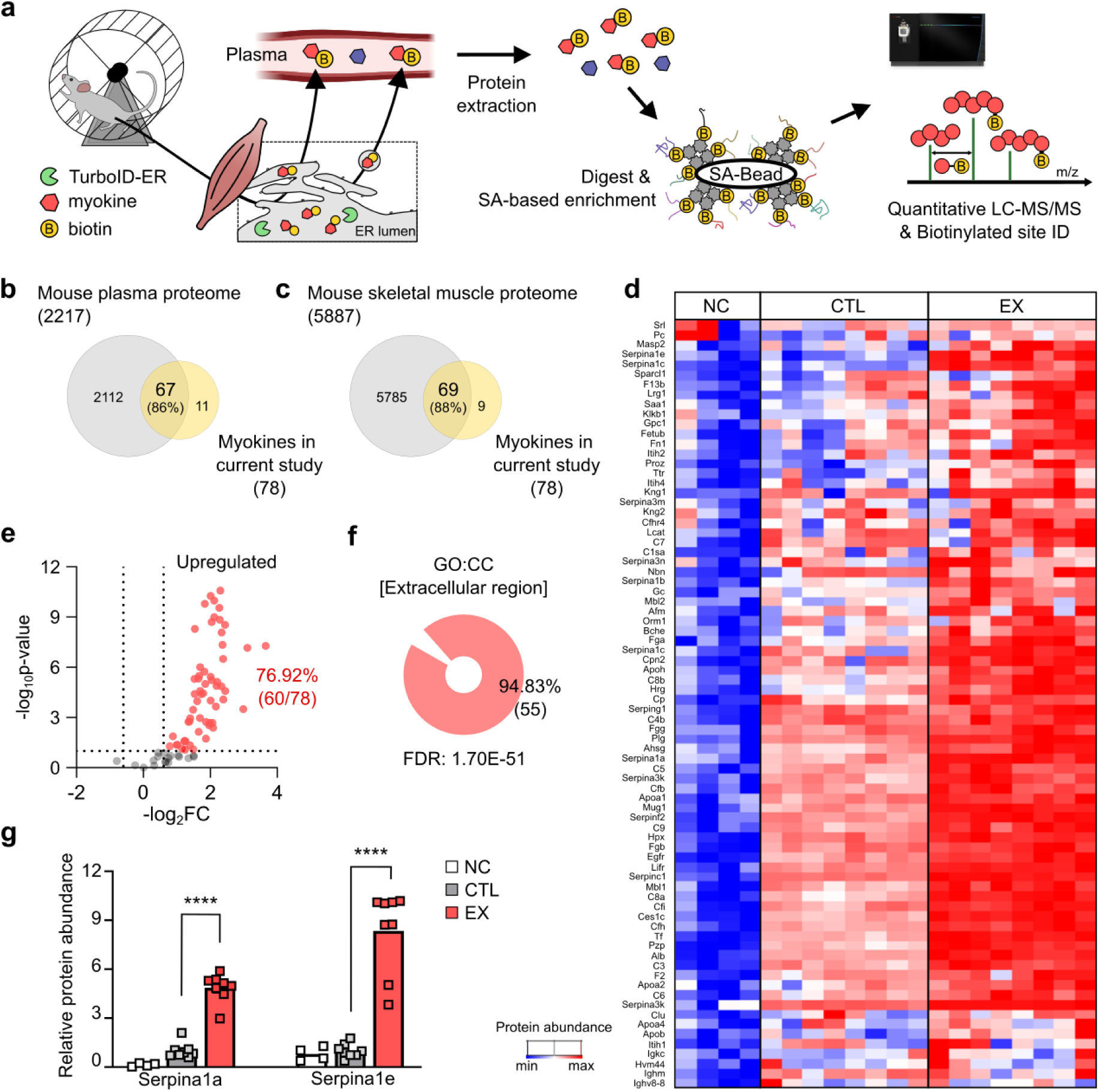
Identification of exercise-induced myokines in plasma. a. Schematic of biotinylated proteins in the plasma of ACTuR mice identified via proteomic profiling. b-c. Venn diagram showing overlap between myokines identified in this study and those identified in a previously reported mouse plasma proteome dataset (b) and a previously reported mouse skeletal muscle proteome dataset (c). d. Heatmap depicting the abundance of plasma myokines. NC: PBS-injected sedentary ACTuR mice; CTL: biotin-injected sedentary ACTuR mice; EX: biotin-injected exercised ACTuR mice. e. Volcano plot illustrating the ∼ 77% (60/78) of myokines whose levels were significantly increased by exercise. (f) GO cellular component (CC) of upregulated myokines postexercise. FDR: false discovery rate calculated by the Benjamini□Hochberg adjustment test. g. Relative protein abundance of Serpina1a and Serpina1e based on mass spectrometry results. ****p<0.0001, one-way ANOVA with Dunnett’s post hoc test. All the data in the bar graphs are presented as the mean ± SEM. Each dot in the graphs represents a tested mouse.

As demonstrated in further analysis, the levels of most of the identified myokines (60 of 78 myokines; ∼ 77%) significantly increased after exercise (p-value< 0.1, −log2(fold change)> 0.6; Extended Data 1; Fig. 2d, e). Gene Ontology (GO) enrichment analysis of upregulated exercise-induced myokines categorized them as cellular components of extracellular region, which coincides with the feature of secreted proteins (Fig. 2f). Furthermore, biological processing terms such as homeostasis, immune response and response to cytokine were enriched from GO term analysis (Fig. 2g), all of which are known to be related to exercise^19,20^. Additional STRING network analysis of enriched putative myokines demonstrated highly interactive networks among these proteins which further suggests these proteins are likely to be involved in similar functional categories (Extended Data Fig. 6c). We found that skeletal muscle-derived Serpina1 family proteins, such as Serpina1a and Serpina1e, were significantly increased in the plasma of exercised ACTuR mice (Fig. 2h). The GO analysis revealed that the term negative regulation of peptidase activity was enriched in these genes, but functions of Serpina1 proteins has yet to be reported to be associated with exercise.

### Serpina1 family proteins are novel exercise-induced myokines

Our proteomics analysis revealed that Serpina1 family proteins are significantly upregulated postexercise, although their functional roles in exercise-induced physiological responses remain unclear. The Serpin family of proteins are abundant in blood and are known to be involved in serine protease inactivation^21^, blood coagulation, and inflammation^22^, as well as being associated with normal liver and lung functions^23,24^. In line with our proteomics results, plasma concentrations of the Serpina1 family were reported to be elevated by 6 weeks of wheel running^1^.

To validate the exercise-induced increase in the amount of muscle-derived Serpina1, we detected the relative amounts of plasma Serpina1 proteins derived from skeletal muscles and non-skeletal muscle tissues (Fig. 3a). The biotinylated Serpina1 proteins were enriched in the plasma fraction of ACTuR mice using streptavidin-conjugated beads (Fig. 3a). The resulting pools of biotinylation-enriched (skeletal muscle-derived) and leftover (non-skeletal muscle-derived) proteins in plasma represented myokines and other organ-derived secretory proteins, respectively. When we assessed the relative amount of Serpina1 protein, we found that the increase in the amount of skeletal muscle-derived Serpina1 in exercised animals was not statistically significant (p= 0.052) but was evident compared with that in sedentary animals (Fig. 3b-e; Extended Data Fig. 7). Owing to the absence of specific antibodies against each mouse Serpina1 family protein, the use of a pan-Serpina1 antibody was the only option for monitoring the plasma level of the Serpina1 protein. Therefore, a statistically nonsignificant increase in plasma Serpina1 protein levels may be due to the mixed responses of Serpina1 family proteins. Despite the limitations of antibody-based differential detection of plasma Serpina1 family proteins, there was no exercise-induced change in the plasma level of non-skeletal muscle-derived Serpina1 family proteins (Fig. 3d; Extended Data Fig. 7), indicating that exercise-induced secretion of Serpina1 proteins from non-skeletal muscle tissues or organs is negligible. Taken together, these results support findings from our proteomic analysis that 4 weeks of exercise increases the level of skeletal muscle-derived Serpina1 in plasma.

**Figure 3.**
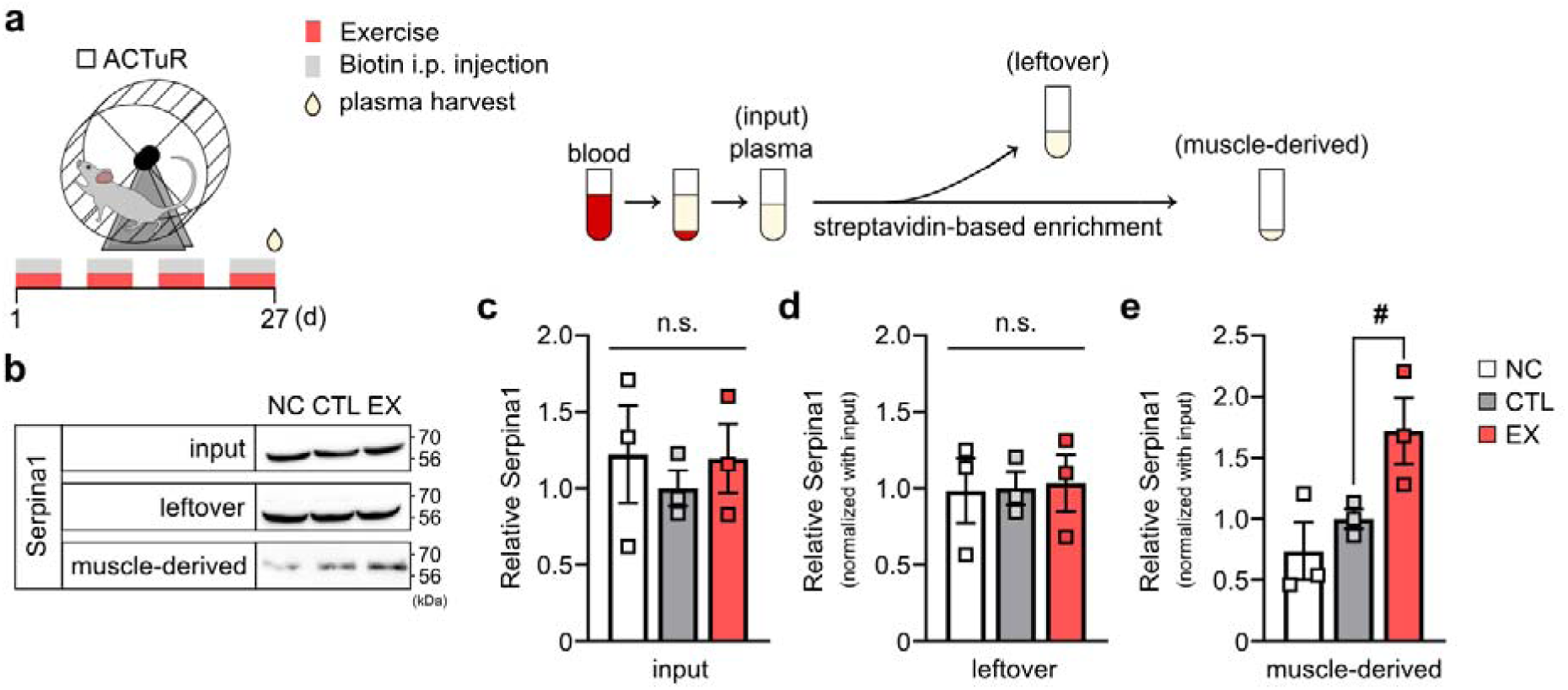
Exercise increases the plasma levels of Serpina1 derived from skeletal muscles. a. Schematic diagram illustrating the process of harvesting exercise-induced plasma proteins (left) and isolating skeletal muscle-derived plasma proteins via enrichment using streptavidin-conjugated beads (right). b. Representative western blot images showing Serpina1 levels in the input, leftover (non-biotinylated proteins remaining after enrichment using streptavidin-conjugated beads), and skeletal muscle-derived (biotinylated proteins enriched/isolated with streptavidin) fractions. NC: PBS-injected sedentary ACTuR mice; CTL: biotin-injected sedentary ACTuR mice; EX: biotin-injected exercised ACTuR mice. c-e. Bar graphs showing average relative Serpina1 levels from the input (c), leftover (d) and skeletal muscle-derived (e) fractions. #p= 0.0520, one-way ANOVA with Sidak’s post hoc test comparing a preselected pair (CTL vs. EX). All the data in the bar graphs are presented as the mean ± SEM. Each dot in the graphs represents a tested mouse.

### Both systemic and muscle-specific overexpression of Serpin1e enhances hippocampal memory

Because the level of skeletal muscle-derived Serpina1 was found to be elevated in the plasma of exercised mice, we hypothesized that exercise-induced memory enhancement is mediated by the direct actions of Serpina1. To test whether Serpina1 itself can mimic the beneficial effects of exercise on learning and memory, V5-tagged recombinant Serpina1 proteins were directly administered to artificially increase the plasma concentration of Serpina1 (Fig. 4, Extended Data Fig. 9, Extended Data Fig. 10). Among the five paralogs of Serpina1 in C57BL/6 mice^24,25^, Serpina1a-d proteins are reported to be highly conserved and expressed at similar levels, with functions of inhibiting the activity of proteases such as elastases^25^. In contrast, Serpina1e is expressed at a lower level than the other Serpina1 paralog proteins and does not have protease inhibitor activity^25^. Accordingly, we selected Serpina1a and Serpina1e for further studies to test their ability to modulate cognition.

**Figure 4.**
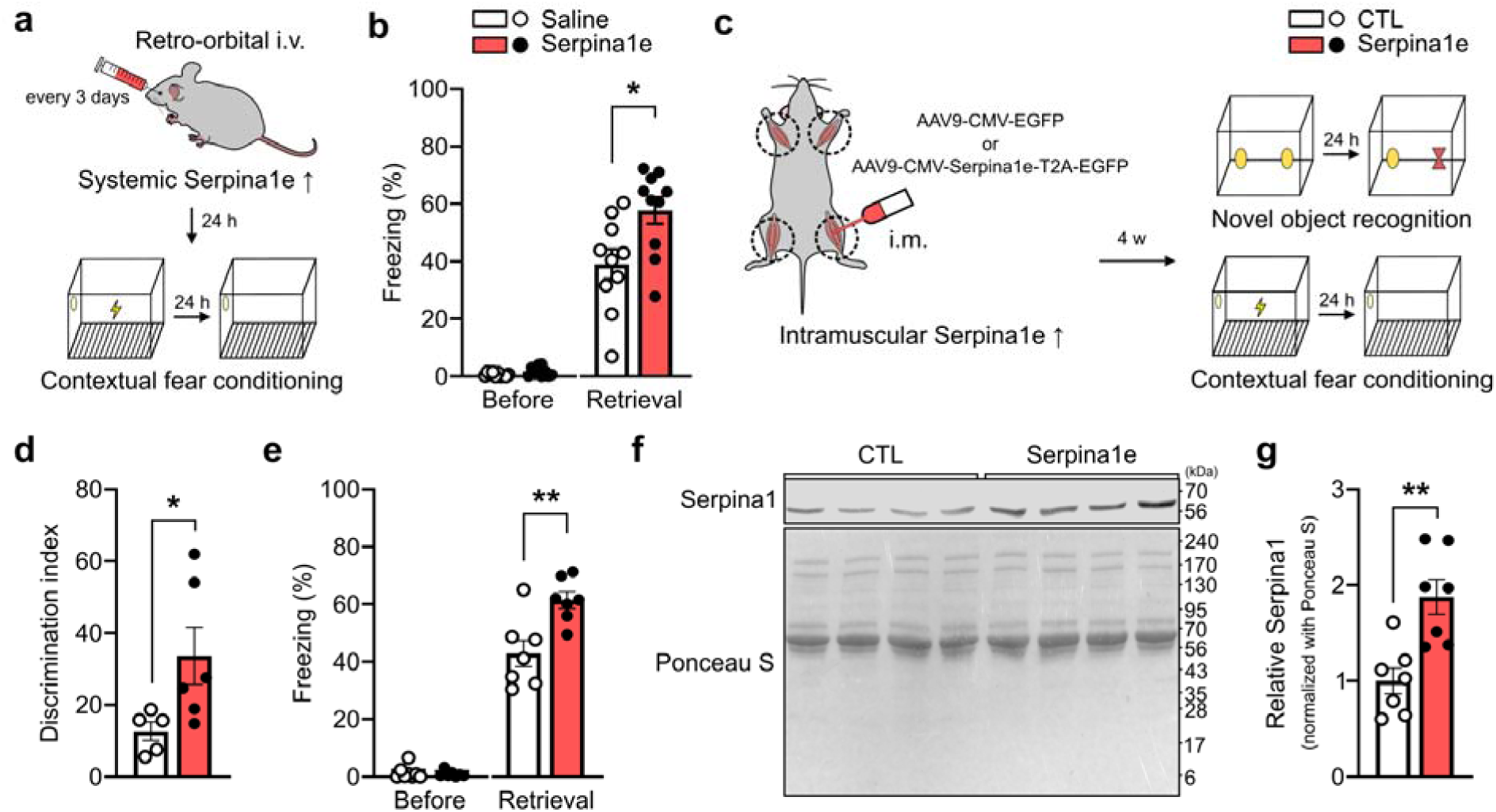
Both systemic and skeletal muscle-specific overexpression of Serpina1e improve hippocampal memory. a. Schematic diagram showing the process of the CFC test after intravenous administration of recombinant Serpina1e-V5. b. Bar graphs depicting the average freezing time (%) in the CFC test. *p= 0.0146, unpaired two-tailed Student’s t test. Saline: PBS i.v. injected mouse; Serpina1e: Serpina1e i.v. injected mouse. c. Schematic diagram showing how the process of hippocampal memory tests is influenced by the overexpression of Serpina1e in skeletal muscle. d. Bar graphs depicting the average discrimination index in the NOR test. *p= 0.0468, unpaired two-tailed Student’s t test. CTL: AAV-CMV-EGFP i.m. injected mouse; Serpina1e: AAV9-CMV-Serpina1e-T2A-EGFP i.m. injected mouse. e. Bar graphs depicting the average freezing time (%) in the CFC test. **p= 0.0049, unpaired two-tailed Student’s t test. f. Representative western blot image showing the levels of plasma Serpina1 (top: Serpina1) and total plasma proteins (bottom: ponceau S) in AAV9-CMV-EGFP (CTL)- or AAV9-CMV-Serpina1e-T2A-EGFP-injected mice (Serpina1e). g. Bar graph depicting the average plasma Serpina1 level (relative to total plasma protein). **p= 0.0022, unpaired two-tailed Student’s t test. All the data in the bar graphs are presented as the mean ± SEM. Each dot in the graphs represents a tested mouse.

To test whether an increase in plasma levels of Serpina1 proteins influences cognitive functions, V5-tagged recombinant Serpina1a and Serpina1e (Serpina1a-V5 and Serpina1e-V5, respectively; 150 µg) were intravenously (i.v.) injected through the retro-orbital vein of wild-type sedentary mice every three days for 4 weeks, while the same volume of saline was injected as a control (Fig. 4a). These systemic administrations of Serpina1a or Serpina1e resulted in no alterations in fear behavior (Fig. 4b, Extended Data Fig. 9b), anxiety levels or general behaviors (Extended Data Fig. 11). Suppression of body weight gain was observed only in the mice injected with Serpina1a (Extended Data Fig. 9c), as was observed in the exercised mice (Extended Data Fig. 5d). This selective effect of Serpina1a on body weight is in line with a previous report that the overexpression of human SERPINA1 alleviates insulin resistance and body weight by inducing an imbalance between the levels of neutrophil elastase and SERPINA1^26^. Sedentary mice administered recombinant Serpina1e exhibited improved performance in the CFC test (Fig. 4b and Extended Data Fig. 8), whereas Serpina1a had no such effect (Extended Data Fig. 9). Hippocampal memory was still improved after systemic coinjection of Serpina1a and Serpina1e (Extended Data Figs. 10 and 11), again supporting the functional role of Serpina1e in memory enhancement.

Next, we validated the beneficial effects of Serpina1e on cognitive functions by testing the memory performance of wild-type mice with skeletal muscle-specific *Serpina1e* overexpression (Fig. 4c). Serpina1e was overexpressed in the skeletal muscles of the four limbs via intramuscular (i.m.) delivery of adeno-associated virus (AAV9) containing CMV promoter-driven *Serpina1e* (Fig. 4c). Our data revealed that sedentary mice with muscular Serpina1e overexpression also exhibited improved performance in the CFC test and novel object recognition test (NOR; Fig. 4c-e), with no significant alterations in locomotion or anxiety levels (Extended Data Fig. 12). Moreover, the plasma level of Serpina1 was significantly elevated by the overexpression of *Serpina1e* in skeletal muscle (Fig. 4f, g). Overall, our results indicate that an increase in the level of plasma Serpina1e leads to enhancements in hippocampus-dependent cognitive functions.

### Plasma Serpina1e can directly migrate to the brain

To dissect the detailed mechanisms by which plasma Serpina1e enhances hippocampal memory, we investigated whether plasma Serpina1e could cross the blood–cerebral spinal fluid (CSF) and brain– blood barriers to reach the brain. We first tested whether plasma recombinant Serpina1e was present in CSF. Pure CSF samples with negligible blood contamination (Extended Data Fig. 13b, c) were collected one hour after intravenous injection of recombinant Serpina1e-V5 (Fig. 5a). We detected recombinant Serpina1e-V5 in not only plasma but also CSF (Fig. 5b), indicating that peripheral Serpina1e can cross the blood–CSF barrier. Moreover, recombinant Serpina1e-V5 was also detected in hippocampal tissues an hour or three hours after intravenous injection (Fig. 5c, d). These results indicate that plasma Serpina1e can cross the blood–CSF and blood–brain barriers, although the underlying mechanisms remain to be elucidated in further studies.

**Figure 5.**
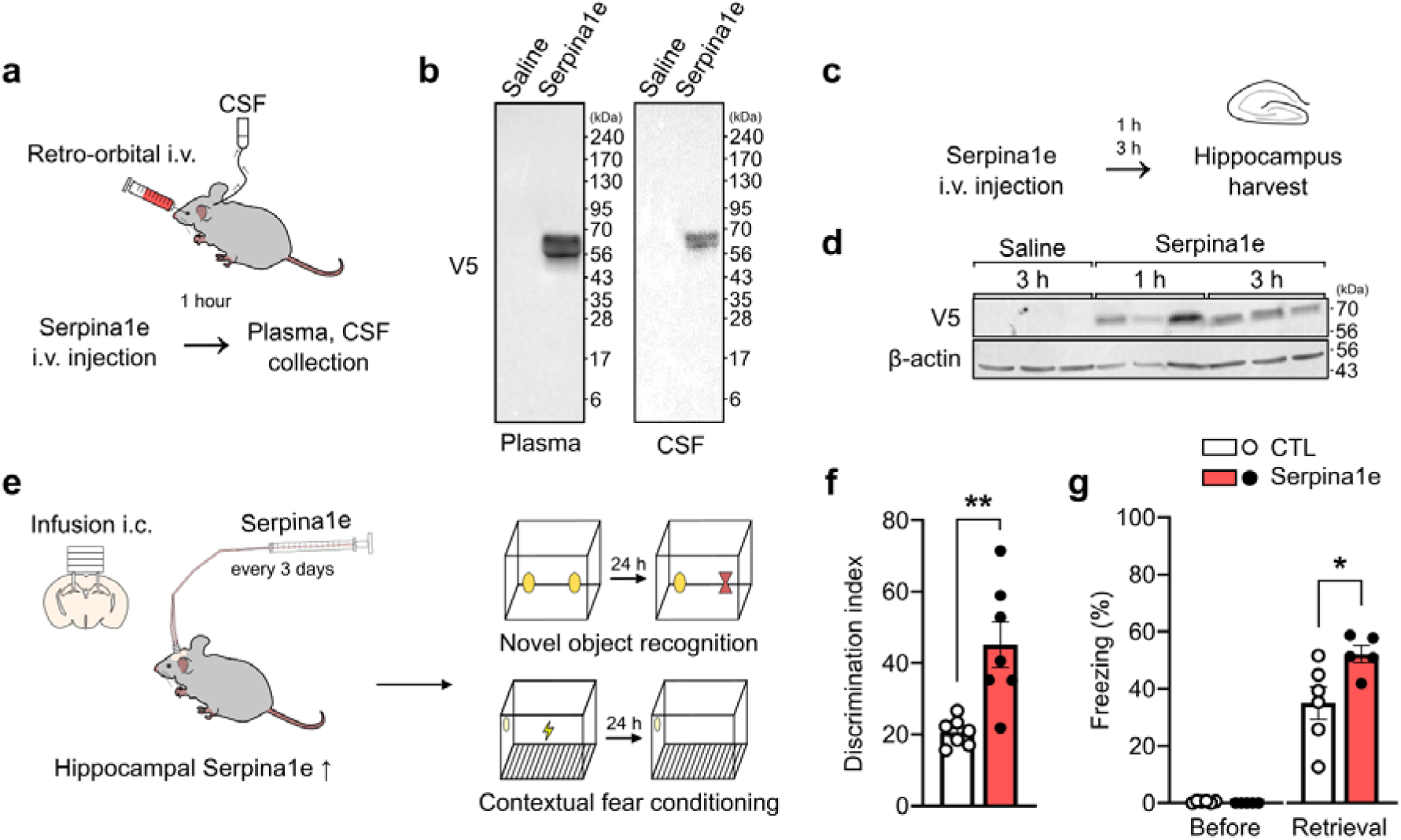
Direct injection of plasma Serpina1e into the hippocampus. a. Schematic diagram showing CSF collection from mice intravenously administered recombinant Serpina1e-V5. b. Representative western blot image showing the detection of recombinant Serpina1e (V5) in plasma and CSF. c. Schematic diagram of the preparation of hippocampal lysates 1 hour or 3 hours after intravenous administration of Serpina1e-V5. d. Representative western blot images showing recombinant Serpina1e (V5) and beta actin (β-actin) in hippocampal lysates. Saline: PBS i.v. injected mouse; Serpina1e: Serpina1e i.v. injected mouse. e. Illustration of the effects of direct infusion of Serpina1e into the hippocampus on cognitive function. f. Bar graphs depicting the average discrimination index in the NOR test. **p=0.0027, unpaired two-tailed Student’s t test. CTL: PBS i.c. infused mouse; Serpina1e: Serpina1e i.c. infused mouse. g. Bar graphs depicting the average freezing time (%) in the CFC test. *p=0.0339, unpaired two-tailed Student’s t test. All the data in the bar graphs are presented as the mean ± SEM. Each dot in the graphs represents a tested mouse.

Given that plasma recombinant Serpina1e can be directly transported to the brain, hippocampal learning and memory may be directly affected by Serpina1e. To test this possibility, Serpina1e (100 ng per hemisphere) was intracerebrally (i.c.) administered into the bilateral dorsal hippocampus of wild-type sedentary mice every three days for 4 weeks (Fig. 5e, Extended Data Fig. 14a). No alterations in body weight gain or general behavior (Extended Data Fig. 14b-d) were observed, similar to the findings in mice intravenously administered Serpina1e (Extended Data Fig. 8c and 11). Twenty-four hours after the last administration, the NOR or CFC test was performed to examine the effects of the hippocampal delivery of Serpina1e on hippocampal cognitive functions (Fig. 5e). Our results revealed that hippocampal memory was significantly enhanced by the hippocampal delivery of recombinant Serpina1e according to both the NOR and CFC tests (Fig. 5f, g), consistent with the effects of the plasma or skeletal muscle-specific overexpression of Serpina1e on hippocampal memory (Fig. 4). Together, these results suggest that the positive impact of exercise on hippocampal function is at least in part mediated by plasma Serpina1e that has directly migrated to the hippocampus.

### Promotion of hippocampal neural growth signaling by Serpina1e

Next, we explored the mechanisms underlying the Serpina1e-induced enhancement of hippocampal learning and memory. Aerobic exercise facilitates synaptic plasticity, which is instrumental in learning and memory^5^, and promotes growth signaling-related molecular and cellular events such as elevated neurogenesis and brain-derived neurotrophic factor (BDNF) upregulation in the brain^27^.

We thus investigated whether Serpina1e can directly induce hippocampal neurogenesis and increase BDNF expression (Fig. 6a). Our results showed that the administration of recombinant Serpina1e to sedentary mice significantly increased the number of 5-bromo-2-deoxyuridine-positive (BrdU^+^) cells in the dentate gyrus (DG) of the hippocampus (Fig. 6b), indicating increased neurogenesis. An increase in BDNF expression was also detected in Serpina1e-injected mice (Fig. 6c). Given that plasma Serpina1e can migrate to the brain (Fig. 5), these results suggest that Serpina1e can directly influence hippocampal neurons and increase neurogenesis and BDNF expression.

**Figure 6.**
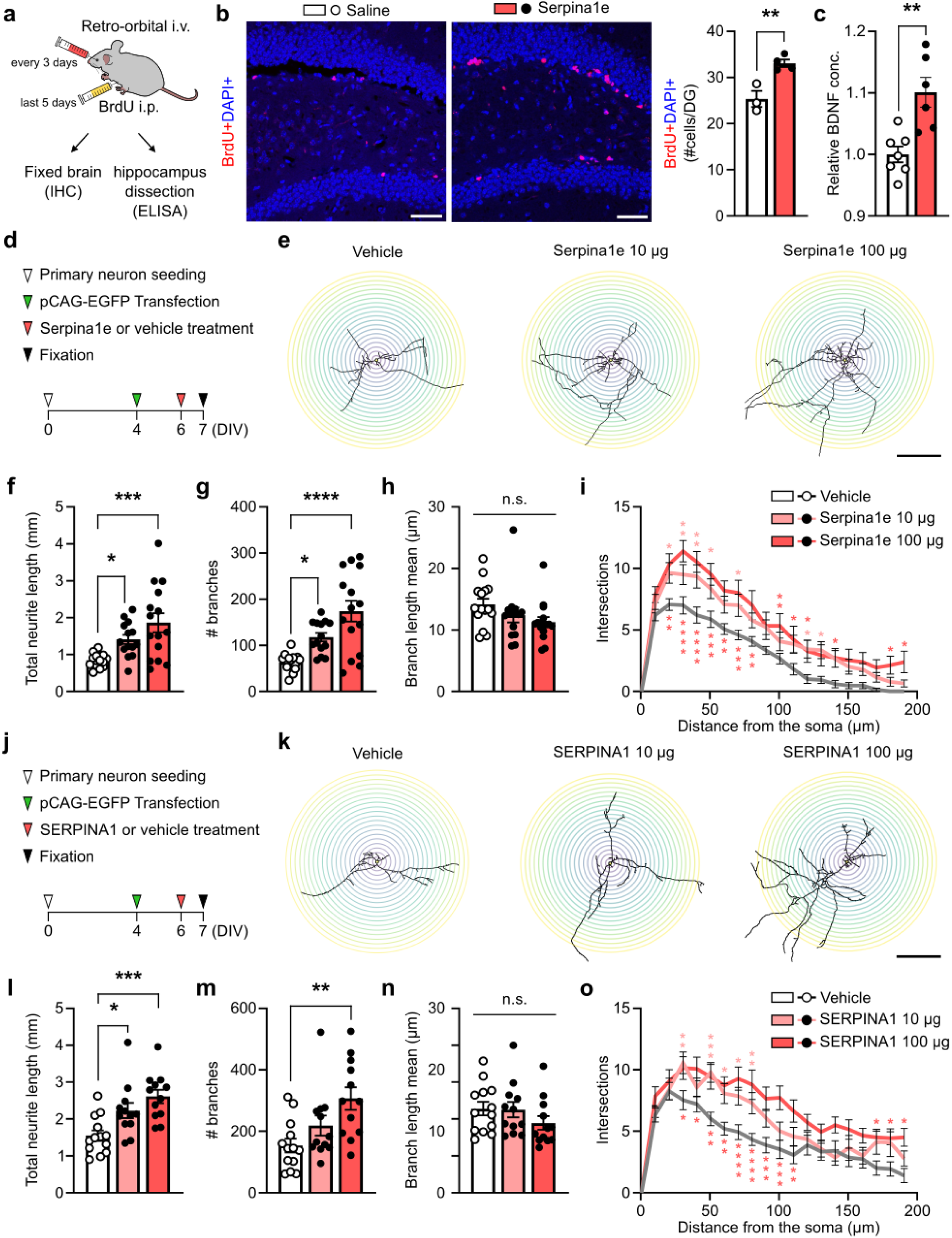
Augmented neuronal growth by Serpina1e. a. Schematic diagram showing intravenous and intraperitoneal injection of recombinant Serpina1e and BrdU, respectively. b. Representative images showing BrdU-positive cells in the hippocampus (left) and bar graphs depicting the average number of BrdU-positive cells in the DG of the hippocampus (right). Scale bar: 50 µm. **p= 0.0066, unpaired two-tailed Student’s t test. Saline: PBS i.v. injected mouse; Serpina1e: Serpina1e i.v. injected mouse. c. Bar graphs depicting relative BDNF levels in the hippocampal lysates. **p= 0.0028, unpaired two-tailed Student’s t test. Each dot in the graphs indicates a tested mouse. d. Experimental timeline for analyzing morphological changes in neurites of cultured primary neurons after Serpina1e application. e. Representative images showing the morphology of cultured neurons treated with vehicle or 10 or 100 µg of Serpina1e. Scale bar: 100 μm. f. Total neurite length. *p= 0.0445, ***p= 0.0002; one-way ANOVA with Dunnett’s post hoc test. g. Total number of branches. *p= 0.0231, ****p<0.0001, one-way ANOVA with Dunnett’s post hoc test. h. Average length of branches. i. Plots depicting the average number of neurite intersections along the distance from the soma. *p< 0.05, **p< 0.01, ***p< 0.001, ****p< 0.0001; two-way ANOVA with Dunnett’s post hoc test. j. Experimental timeline for analyzing morphological changes in the neurites of cultured primary neurons after SERPINA1 application. k. Representative images showing the morphology of cultured neurons treated with vehicle or 10 or 100 µg of SERPINA1. Scale bar: 100 μm. l. Total neurite length. *p= 0.0186, ***P= 0.0003; one-way ANOVA with Dunnett’s post hoc test. m. Total number of branches. **p= 0.0033, one-way ANOVA with Dunnett’s post hoc test. n. Average length of branches. o. Plots depicting the average number of neurite intersections along the distance from the soma. *p< 0.05, **p< 0.01, ***p< 0.001, ****p< 0.0001; two-way ANOVA with Dunnett’s post hoc test. All data are presented as the mean ± SEM. Each dot in the graphs represents a tested cultured neuron at least from three different culture batches.

Neurotrophic factors, such as BDNF, promote neurite growth and increase the morphological complexity of neurons^28^. To test whether Serpina1e can directly induce such changes in neurons, we treated primary cultured neurons with recombinant Serpina1e (Fig. 6d). Our data revealed that Serpina1e-treated neurons showed increases in neurite length, branch number (Fig. 6e-g), and complexity (Fig. 6i). These effects were dependent on the concentration of Serpina1e because the Serpina1e-induced changes in neurite structure were proportional to the number of recombinant Serpina1e-treated cultured neurons (Fig. 6e-g and 6i). Similar positive effects of Serpina1e on neurites were also found in cultured neurons treated with human SERPINA1 (Fig. 6j-o), suggesting that the neuron growth function of SERPINA1 is conserved in Serpina1e. Because recombinant SERPINA1 has an anti-inflammatory function independent of its anti-protease activity^28^, Serpina1e- and SERPINA1-induced structural growth in neurons is suggested to be mediated by the upregulation of growth signals, such as BDNF, via anti-inflammatory effects^28,29^.

## Discussion

Our study identified Serpina1e as a novel molecular mediator of interactions between the peripheral and central nervous systems, playing a key role in exercise-induced cognitive improvements. Despite previous reports showing an increase in the plasma levels of mouse Serpina1 and human SERPINA1 after exercise^1,10^, our study is the first to reveal that it is derived from skeletal muscles. We exploited ACTuR mice, which allows the selective labeling of skeletal muscle-originated secretory proteins with biotin, to obtain molecular information on the muscle secretome postexercise. Regardless of its high expression in and secretion from the liver^30^, the ACTuR model allowed us to detect skeletal muscle-derived Serpina1 and discriminate skeletal muscle-derived Serpina1 from non-skeletal muscle-derived Serpina1 in the plasma of exercised animals (Figs. 2 and 3). Our data suggest that Serpina1 is not only a liver-derived factor but also an exercise-induced myokine.

An increased amount of plasma Serina1 may influence diverse physiological functions^31,32^, and our study provides evidence that an increase in systemic Serpina1 levels is sufficient for enhancing hippocampal learning and memory (Figs. 4 and 5). The mechanisms underlying Serpina1e-induced cognitive enhancements are not fully understood, but we provide evidence for the upregulation of neural growth factors by Serpina1e delivered to the brain (Fig. 6), possibly via the immunomodulatory or anti-inflammatory actions of Serpina1e. In fact, exercise is known to increase the plasma levels of anti-inflammatory peripheral proteins such as Clusterin (Fig. 2d, Extended Data 1; ref. 10), and the migration of such proteins across several physiological barriers to the brain could reduce neuroinflammatory gene expression^10^. Serpina1e was reported to have no anti-protease activity, possibly due to a change in the P1 amino acid in the reactive center loop (RCL), but could possess anti-inflammatory properties independent of its anti-protease activity, similar to SERPINA1^29,33^. Therefore, reduced neuroinflammation caused by Serpina1e is likely associated with the upregulation of neuronal growth factors, such as BDNF (Fig. 6), resulting in increased neurite growth and neurogenesis. However, mechanisms other than anti-inflammatory or BDNF-dependent signaling are likely also involved.

Overall, this study presents a novel myokine, Serpina1e, which we propose is a key mediator of exercise-induced cognitive enhancements. Our findings offer new strategies for developing therapies to ameliorate cognitive dysfunction caused by normal aging or neurodegenerative diseases.

## Acknowledgment

This work was supported by grants from the National Research Foundation of Korea (NRF) funded by the Korean government (MSICT) [NRF-2021R1A2C109399112 to H.P., RS-2024-00343424 to J.-S.K.] and Institute for Basic Science from the Ministry of Science and ICT of Korea [IBS-R008-D1 to J.-S.K.]. Data from confocal microscope imaging and behavior tests were acquired using instruments at the Brain Research Core Facilities of the Korea Brain Research Institute.

## Author contributions

H.K. and H.P. designed the projects. H.K. performed all behavioral and biochemical experiments with ACTuR and WT mice and recombinant Serpina1 proteins. S.S. and J.-S.K. performed the mass spectrometry experiments and proteomic analyses of biotinylated plasma proteins. J.H. performed experiments for detecting recombinant Serpina1 proteins in CSF and evaluating the effects of recombinant Serpina1 and SERPINA1 proteins on the neurite morphology of cultured neurons. H.P. supervised the project. H.P., H.K., S.S., and J.-S.K. wrote the paper.

## Methods

### Animals

The following mouse lines were used: C57BL/6 mice (The Jackson Laboratory), ACTA1-Cre mice (The Jackson Laboratory), LSL-TurboID-ER (loxP-stop-loxP-TurboID-V5-KDEL generated from Cyagen) mice, and ACTuR (ACTA1-Cre:LSL-TurboID-ER) mice. ROSA26 was targeted by a dual-guide RNA (GGCAGGCTTAAAGGCTAACCTGG, CTCCAGTCTTTCTAGAAGATGGG) for the insertion of the TurboID-ER coding sequence via CRISPR/Cas9-mediated editing to generate LSL-TurboID-ER mice. To label myokines and identify exercise-induced myokines, the *Acta1*-Cre and LSL-TurboID-ER lines were crossed to produce ACTuR (*Acta1*-Cre:LSL-TurboID-ER) mice. All the mouse lines were maintained on a C57BL/6 genetic background. The mice were housed in a specific pathogen-free room at the Laboratory Animal Center in the Korea Brain Research Institute (KBRI) that was kept at 20–24°C and 40–60% humidity under a 12–12 h light–dark cycle with dark hours between 20:00–08:00. All mouse experiments were performed in accordance with all ethical regulations and institutional guidelines approved by the Institutional Animal Care and Use Committee (IACUC) of the KBRI. Young male adult (8–11-week-old) mice were randomly assigned to different groups for the experiments. For western blot analysis and LC□MS/MS with myokines in plasma, ACTuR mice were used. For behavioral analysis after protein injections, protein infusions or viral infections, C57BL/6 wild type mice were used.

### Behavioral experiments

Behavioral testing of sedentary or exercised mice began the day after the last day of exercise. Behavioral testing of mice intravenously injected with Serpina1a or Serpina1e for 4 weeks began the day after the final injection. Also, behavioral testing of mice that received multiple infusions of Serpina1e into the hippocampus began the day after the final infusion. Additionally, behavioral testing of mice intramuscularly injected AAV into all four limbs began after 4 weeks from the injection day. The experimenters were blinded to the treatment during testing and analysis.

#### Voluntary exercise

Male mice (aged 8 to 11 weeks) were housed in pairs either in activity cages with access to an activity wheel or in standard housing cages (controls) for 4 weeks. All exercise experiments were performed in activity cages with a running wheel installed. The total running distance was calculated based on the number of wheel revolutions recorded by a revolution counter. Activity cages with a running wheel and revolution counter were purchased from Ugo Basile. Exercised mice were housed in the activity cages in pairs with free access to unlocked wheels at night (20:00-08:00) by following a 5-day-per-week regimen for four weeks. The control mice were housed in pairs in standard animal facility cages.

#### Open field test

The open field test was used to determine general activity levels and gross locomotor activity. The assessment took place in a square arena within a specially designed sound attenuating chamber. The arena was 30 cm (L) × 30 cm (W) × 25 cm (H). The mouse was placed in the corner of the testing arena and allowed to explore the arena for 5 min while being tracked by an automated tracking system of SMART3.0 (Panlab) software. Parameters, including distance moved, velocity, and time spent in predefined zones of the arena, were recorded. The arena was cleaned with distilled water and 70% ethanol at the end of the trial.

#### Elevated plus maze (EPM) test

The customized EPM apparatuses had two open arms measuring 30 cm in height and 6 cm in width and two closed arms with a maze wall height of 20 cm and 45 cm in height from the floor. The mice started the EPM test at the center of the maze and freely explored for 5 min while the video was recorded. Parameters, including distance moved and time spent in predefined zones of the arena, were recorded and analyzed with SMART 3.0 (Panlab) software. The arena was cleaned with distilled water and 70% ethanol at the end of the trial.

#### Forced swimming test

The forced swimming test was performed in a plastic cylinder (26 cm height, 11 cm diameter) filled to a depth of 18.5 cm with water (24–25°C). The mice were exposed to the plastic cylinder for 15 min on the first day. After 24 h, the mice were placed into the same plastic cylinder for 5 min, and the data were recorded by a video camera. The immobility time and latency to the first bout of immobility were manually evaluated.

#### Rotarod test

The accelerated rotarod test was performed on a rotarod (Ugo Basile) with two sessions at 1 h intervals. Each session was subjected to three trials for 5 min with an acceleration of 4–40 rpm, and the results were recorded. For the analysis, the trial showing the longest latency to fall from the second session was selected as the indicator of motor coordination.

#### Morris water maze test

The Morris water maze was conducted in a circular pool (120 cm diameter). Tempera paint was added to the water until it became opaque, and a hidden platform (17 cm diameter) was placed 1 cm below the water surface. The water temperature was maintained at ±1°C from 21°C. Black curtains surrounded the water tank, and distinct visual cues were hung from the curtains. The mice were monitored via the EthoVision XT (Noldus Information Technology) video tracking system directly above the water tank, and the parameters were measured using EthoVision software. Pseudorandomized platform locations and drop locations were recorded for each mouse, with one trial lasting 60 sec. The trial ended either when the subject rested on the hidden platform for 5 s or when the duration of the trial expired. The trial was repeated four times for each mouse during each day of training. Five days of training were performed, and two days of probe trials were conducted following training. The time to reach the platform was recorded to obtain a learning curve graph of latency to escape. The time spent in the target quadrant, swimming speed, total distance moved and number of platform crossings were analyzed.

#### CFC tests

CFC tests were performed in sound-attenuating chambers equipped with steel shocking floors (Panlab). Freezing behavior in the cages was recorded by a low-light camera, and the shock and freezing behavior analyses were automatically conducted by Packwin software (Panlab). During fear-conditioning training, the mice were habituated to the shock cage for 3 min and then subjected to pairing of a context (box) with a single foot shock for 2 sec at 0.5 mA. The mice were left in the shock cage for an additional 1 min after the shock and then returned to their home cage. The next day, the mice underwent contextual recall testing; contextual testing was performed in the same cages that were used during the training trial. Mouse freezing behavior was tracked over a period of 3 min. To examine long-term memory, the mice were subjected to contextual recall testing in the same context three weeks after the training day.

#### NOR test

NOR tests were conducted in the same arena where the open field tests were previously carried out. On the first day, the mice were habituated to the empty arena for 5 min. After habituation, a pair of identical objects were positioned at distinct corners of the arena, and subsequently, the mice were given a 5-min exploration period within the arena. This procedure facilitated the establishment of baseline familiarity with the objects. After 24 hours, the mice were introduced to a familiar object or a novel object within the same area to assess their discriminatory ability. The discrimination index (DI) was calculated using the time spent exploring a novel object and the time spent exploring a familiar object. The formula was as follows:

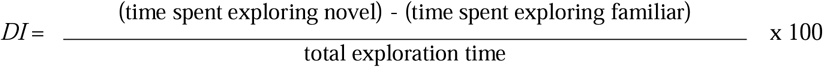

### Preparation of biotinylated myokines in plasma and tissues

To induce the biotinylation of myokines in ACTuR mice, we intraperitoneally injected biotin (24 mg/kg) dissolved in phosphate-buffered saline (PBS, Gibco). ACTuR mice were euthanized between 08:30 and 10:00. The mice were anesthetized with 1 ml of isoflurane (Piramal) in a sealed plastic box in preparation for perfusion or plasma collection. Blood was collected from the right heart ventricle with 5 µl of 0.5 M EDTA (Sigma□Aldrich) and centrifuged at 4°C for 20 min at 12,000 rpm to collect the plasma. A protease inhibitor cocktail (P.I.C., Quartett) was added to the plasma. Tissues (brain, skeletal muscle, liver, lung, and heart) were harvested and homogenized with a bead homogenizer (Taco) and RIPA lysis buffer (Rockland). The tissue lysates were harvested from the supernatants of the tissue samples following centrifugation at 4°C for 20 min at 12,000 rpm.

### Western blot analysis

The protein concentration was determined using a BCA assay kit (Thermo Scientific) according to the manufacturer’s instructions. The samples were boiled for 3 min at 95°C in Laemmli buffer (Bio-Rad) and separated on precast 4□20% polyacrylamide gels (Bio-Rad). Afterward, the proteins were transferred onto 0.2 µm PVDF membranes via a Turbo transfer system (Bio-Rad). The membranes were blocked for 1 h in 3% skim milk (Sigma□Aldrich) in 1x TBS (Bio-Rad) with 0.1% Tween 20 (Sigma□Aldrich), incubated overnight at 4°C with the primary antibody, incubated for 2 h at room temperature with the HRP-conjugated secondary antibody and developed using an ECL prime solution (Thermo Scientific). The antibodies used were rabbit anti-V5 (Cell Signaling Technology), rabbit anti-Serpina1 (Invitrogen), HRP-conjugated anti-rabbit IgG (Bio-Rad), streptavidin-HRP (Thermo Scientific) and rabbit anti-β-actin-HRP antibodies (Cell Signaling Technology).

### Mass spectrometry

#### Sample preparation

Harvested plasma samples were lysed with 2% SDS in 1× TBS (25 mM Tris, 0.15 M NaCl, pH 7.2; Thermo Scientific) supplemented with a 1 × protease inhibitor cocktail. The lysates were clarified via ultrasonication (Bioruptor, diagenode) for 15 min in a cold-water bath. To remove free probes, 6 times the sample volume of cold acetone (−20°C, Sigma□Aldrich) was added to each lysate and kept at −20°C. After at least two hours, the samples were centrifuged at 15,000 × g for 15 min at 4°C. The supernatant was gently removed, and acetone precipitation was repeated with 5 mL of cold acetone containing a □ volume of 1× TBS. After the supernatant was removed, the pellet was solubilized with 8 M urea (Sigma□Aldrich) in 50 mM ammonium bicarbonate (ABC, Sigma□Aldrich). The protein concentration was measured, and the samples were denatured at 650 rpm for 1 h at 37°C. After denaturation, the samples were reduced with a final concentration of 10 mM dithiothreitol (Sigma□Aldrich) and incubated at 650 rpm for 1 h at 37°C. The samples were alkylated by adding 40 mM iodoacetamide (Sigma□Aldrich) to the final concentration and mixing at 650 rpm for 1 h at 37°C. The samples were diluted eight times with 50 mM ABC, and CaCl2 (Alfa Aesar) was added to achieve a final concentration of 1 mM. Approximately 2 mg of protein sample was digested by adding trypsin (Thermo Scientific, 50:1 w/w) and incubated at 650 rpm for 6–18 h at 37°C. After trypsinization, the digested peptide samples were incubated with 150 µL of streptavidin beads (Pierce, 88817), which were prewashed with 2 M urea in 1× TBS four times prior to sample enrichment. The samples were incubated for 1 h at room temperature with end-over-end rotation. To remove nonspecifically bound peptides, the beads were washed twice with 2 M urea in 50 mM ABC and then finally washed with distilled water. To elute the biotinylated peptides, elution buffer [80% acetonitrile (Sigma□Aldrich), 0.2% TFA (Sigma□Aldrich), and 0.1% formic acid (Thermo Scientific)] was added, and the mixture was incubated at 60°C for 5 min. Each supernatant was transferred to new tubes, and the elution step was repeated 2-3 times. The combined elution fractions were dried using a SpeedVac (Eppendorf). The samples were stored at −20°C or injected for mass spectrometry directly.

#### LCJMS/MS analysis of enriched peptide samples

Analytical capillary columns (100 cm × 75 µm i.d.) and trap columns (2 cm × 150 µm i.d.) were packed in-house with 3 µm Jupiter C18 particles (Phenomenex, Torrance). The long analytical column was placed in a column heater (Analytical Sales and Services) regulated to a temperature of 45°C. The NanoAcquity UPLC system (Waters, Milford) was operated at a flow rate of 300 nL/min over 2 h with a linear gradient ranging from 95% solvent A (H2O with 0.1% formic acid) to 40% solvent B (acetonitrile with 0.1% formic acid). The enriched samples were analyzed on an Orbitrap Fusion Lumos mass spectrometer (Thermo Scientific) equipped with an in-house customized nanoelectrospray ion source. Precursor ions were acquired (m/z 300-1500) at 120 K resolving power, and the precursor for MS/MS analysis was isolated at 1.4 Th. Higher-energy collisional dissociation (HCD) with 30% collision energy was used for sequencing with an autogel gain control (AGC) target of 1e^5^. The resolving power for the acquired MS2 spectra was set to 30k with a maximum injection time of 200 ms.

### Proteomics

#### MS data processing and protein identification

All MS/MS data were searched by MaxQuant (version 1.6.2.3) with the Andromeda search engine at 10 ppm precursor ion mass tolerance against the *Mus musculus* proteome database (UP000000589, 55,367 entries, UniProt (www.uniprot.org), 2021-06-22). Label-free quantification (LFQ) and match-between runs were used with the following search parameters for the TurboID experiment: partial trypsin digestion, fixed carbaminomethylation on cysteine, dynamic oxidation of methionine, protein N-terminal acetylation with biotin (+226.0775) labels of lysine residues. A false discovery rate (FDR) of less than 1% was obtained for unique labeled peptides and unique labeled proteins. LFQ intensity values were log-transformed for further analysis, and missing values were filled by imputed values representing a normal distribution around the detection limit. To impute the missing values, first, the intensity distributions of the mean and standard deviation were determined; then, for the imputation values, a new distribution based on a Gaussian distribution with a downshift of 1.8 and width of 0.3 standard deviations was created for the total matrix.

#### Principal component analysis (PCA) and GO enrichment analysis

Perseus (ver. 1.6.2.3) raw values were used to perform the PCA, and each set of experiments was normalized to the interval (0-2) independently and subjected to hierarchical clustering analysis with the following options: Euclidean distance with the option of preprocessing with k-means clustering, 10 maximal number of iterations and preserving order of column constraints. Defined genes within each cluster were subjected to GO enrichment analysis via the DAVID online tool (https://david.ncifcrf.gov/). The background used for GO analysis for each cluster was set as the entire proteome of *Homo sapiens*.

### Intravenous administration of Serpina1a and Serpina1e

Recombinant Serpina1a and Serpina1e with V5 (GKPIPNPLLGLDST) and His tags (HHHHHH)were synthesized with amino acid (aa) sequences as following:

-Serpina1e-V5 (433 aa)
MTPSISWCLLLLAGLCCLVPSFLAEDVQETDTSQKDQSPASHEIATNLGDFAISLYRELVHQSNTSNIFFSPVSIATAFAMLSLGSKGDTHTQILEGLQFNLTQTSEADIHNSFQHLLQTLNRPDSELQLSTGNGLFVNNDLKLVEKFLEEAKNHYQAEVFSVNFAESEEAKKVINDFVEKGTQGKIVEAVKKLEQDTVFVLANYILFKGKWKKPFDPENTKQAEFHVDESTTVKVPMMTLSGMLDVHHCSTLSSWVLLMDYAGNATAVFLLPDDGKMQHLEQTLNKELISKFLLNRRRRLAQIHIPRLSISGNYNLETLMSPLGITRIFNSGADLSGITEENAPLKLSQAVHKAVLTIDETGTEAAAATVLQGGFLSMPPILHFNRPFLFIIFEEHSQSPLFVGKVVDPTHKGKPIPNPLLGLDSTHHHHHH
-Serpina1a-V5 (433 aa)
MTPSISWGLLLLAGLCCLVPSFLAEDVQETDTSQKDQSPASHEIATNLGDFAISLYRELVHQSNTSNIFFSPVSIATAFAMLSLGSKGDTHTQILEGLQFNLTQTSEADIHKSFQHLLQTLNRPDSELQLSTGNGLFVNNDLKLVEKFLEEAKNHYQAEVFSVNFAESEEAKKVINDFVEKGTQGKIAEAVKKLDQDTVFALANYILFKGKWKKPFDPENTEEAEFHVDESTTVKVPMMTLSGMLHVHHCSTLSSWVLLMDYAGNATAVFLLPDDGKMQHLEQTLSKELISKFLLNRRRRLAQIHFPRLSISGEYNLKTLMSPLGITRIFNNGADLSGITEENAPLKLSQAVHKAVLTIDETGTEAAAVTVLQMVPMSMPPILRFDHPFLFIIFEEHTQSPIFLGKVVDPTHKGKPIPNPLLGLDSTHHHHHH

Recombinant Serpina1a-V5 or Serpina1e-V5 (150 µg/100 µl) were retro-orbitally injected into sedentary C57BL/6 mice for 4 weeks every three days, for a total of 9 times. For the Serpina1 transmission experiments, sedentary mice were injected retro-orbitally with Serpina1e-V5 at 300 µg/100 µl and anesthetized to collect CSF and hippocampal tissues.

### Intramuscular injection of AAV9

To investigate whether muscular Serpina1e overexpression induces cognitive enhancement, we delivered a single dose of AAV9-CMV-Serpina1e-T2A-EGFP (VectorBuilder; 1.06 × 10e^12^ GCs/mouse) into each of the four limbs of C57BL/6 mice. The control mice were injected with the same dose of AAV9-CMV-EGFP (VectorBuilder; 1.07 × 10e^12^ GC/mouse). Behavioral assessments were performed four weeks after viral infection.

### Infusion of recombinant Serpina1e into the hippocampus

To address the direct effects of Serpina1e treatment on the hippocampus, we infused Serpina1e-V5 into the the hippocampus (AP −1.95, ML ±1.20, and DV −1.70; Extended Data Fig. 14e) with the same timeline as systemic injections. A custom dual-guide cannula was designed with a 26-gauge, 7.8 mm pedestal height, 2.4 mm distance and 2 mm distance below the pedestal (RWD). The dual-guide cannula was implanted and covered with a dummy cannula with a 1 mm projection to fit the dual-guide cannula. A total of 100 ng/250 nl of Serpina1e per hemisphere was infused through the implanted cannula for 4 weeks every 3 days at 100 nl/min. The actual infused site in the hippocampus was determined by injecting DiI dye through the implanted cannula when all behavioral tests had concluded.

### BrdU labeling

To examine the effects of Serpina1e on hippocampal neurogenesis, 5-bromo-2′-deoxyuridine (BrdU, Sigma□Aldrich) was injected intraperitoneally at 50 mg/kg into sedentary mice. The BrdU stock was resuspended in PBS at 50 mg/ml and stored at −20°C. The control mice were injected with an equivalent volume of PBS.

To image BrdU-positive cells in the hippocampus, the mice were anesthetized as previously described and transcranially perfused with 4 % paraformaldehyde (PFA, Biosesang) following perfusion with 1x PBS. The brain tissue was stored in 4% PFA for postfixation overnight and then switched to 30% sucrose solution for an additional 48 h in preparation for brain sectioning. Serial coronal sections of the hippocampus (thickness, 40 µm) were collected using a cryostat (Leica, CM3050S) and washed with PBS. All brain sections were blocked with blocking solution (10% normal goat serum, 0.2% Triton X-100 in PBS) for 1 h at room temperature and then incubated in primary antibody solution. All brain sections stained for BrdU were pretreated with 2 M HCl for 30 min at 37°C before blocking. The primary antibody used for immunostaining was a rat anti-BrdU antibody (1:2,500, Abcam), and the fluorescent secondary antibodies were diluted with blocking solution at a concentration of 1:200 and incubated with the sections for 2 h at room temperature. The fluorescent secondary antibody used was a goat anti-rat IgG Alexa Fluor 568-conjugated antibody (Thermo Scientific). Nuclei were fluorescently labeled with DAPI containing mounting solution (Vectashield).

### Enzyme-linked immunosorbent assays of BDNF

To measure BDNF levels in the hippocampus, C57BL/6 mice were intravenously injected with Serpina1e for 4 weeks every 3 days and then sacrificed 1 hour after the last injection. Hippocampal tissues were harvested following perfusion with PBS and lysed with lysis buffer (1% Triton X-100, 0.1% Tween 20, 100x P.I.C., and 1x PBS). The BDNF concentration was determined via enzyme-linked immunosorbent assays (Novusbio) according to the manufacturer’s instructions.

### Primary cortical neuron culture

Cortical neurons were prepared from the brains of C57BL/6 embryos at embryonic days 16-17. Cortices were dissected in dissection medium (10 mM HEPES in HBSS, Gibco), incubated in 0.25% trypsin-EDTA (Gibco) in a 37°C water bath for 20 min, and then triturated with 2 ml of plating medium (2 mM L-glutamine (Gibco), 0.45% glucose (Sigma□Aldrich), 10% FBS (Thermo Scientific), 5,000 U/ml penicillin and 5,000 µg/ml streptomycin in MEM (Gibco)). Dissociated cells were filtered through a cell strainer and plated on poly-L-lysine-coated coverslips in 12-well plates (3 × 10^5^ cells/well) in plating media. After one hour, the plates were replaced with maintenance medium (2% B27 supplement, Thermo Scientific), 2 mM L-glutamine, 5,000 U/ml penicillin and 5,000 µg/ml streptomycin in neurobasal medium (Gibco)).

### Acquisition and analysis of fluorescence images

Immunofluorescence images were obtained, quantified, and analyzed by an observer who was blinded to the treatment. For neurogenesis analysis via BrdU staining, fluorescence images were acquired by using a STELLARIS 8 microscope (Leica, 40x/1.10 NA) with the pinhole set to 1 Airy unit. Images were taken at the suggested optimal value interval based on magnification and analyzed using ImageJ (National Institutes of Health). The number of BrdU-labeled cells was manually counted in 12 sections per mouse. To analyze neurite morphology, cortical neurons at DIV7 were used. The neurons were imaged using a Leica SP8 confocal microscope (20x/0.75 NA). The images were processed for skeletonization via ImageJ/FIJI, and the total neurite length, branch number, and branch length were analyzed. The skeletonized images were further analyzed via the Sholl analysis plugin in ImageJ/FIJI, which calculates the number of neurite crossings at intervals of 10 µm from the cell body.

### Statistics

All the statistical analyses were performed via GraphPad Prism 8 with 95% confidence, and all the values are expressed as the mean ±SEM. One-way ANOVA with Dunnett’s or Sidak’s post hoc tests was used to compare three groups, and unpaired two-tailed Student’s t tests were used to compare two groups. The data in Fig. 6i and 6o were analyzed via two-way ANOVA with Dunnett’s post hoc test. The statistical tests for each experiment are provided in the legends of the respective figures. P values less than 0.05 were considered statistically significant.

