## Extended Data Fig. for "Serpina1e mediates the exercise-induced enhancement of hippocampal memory"

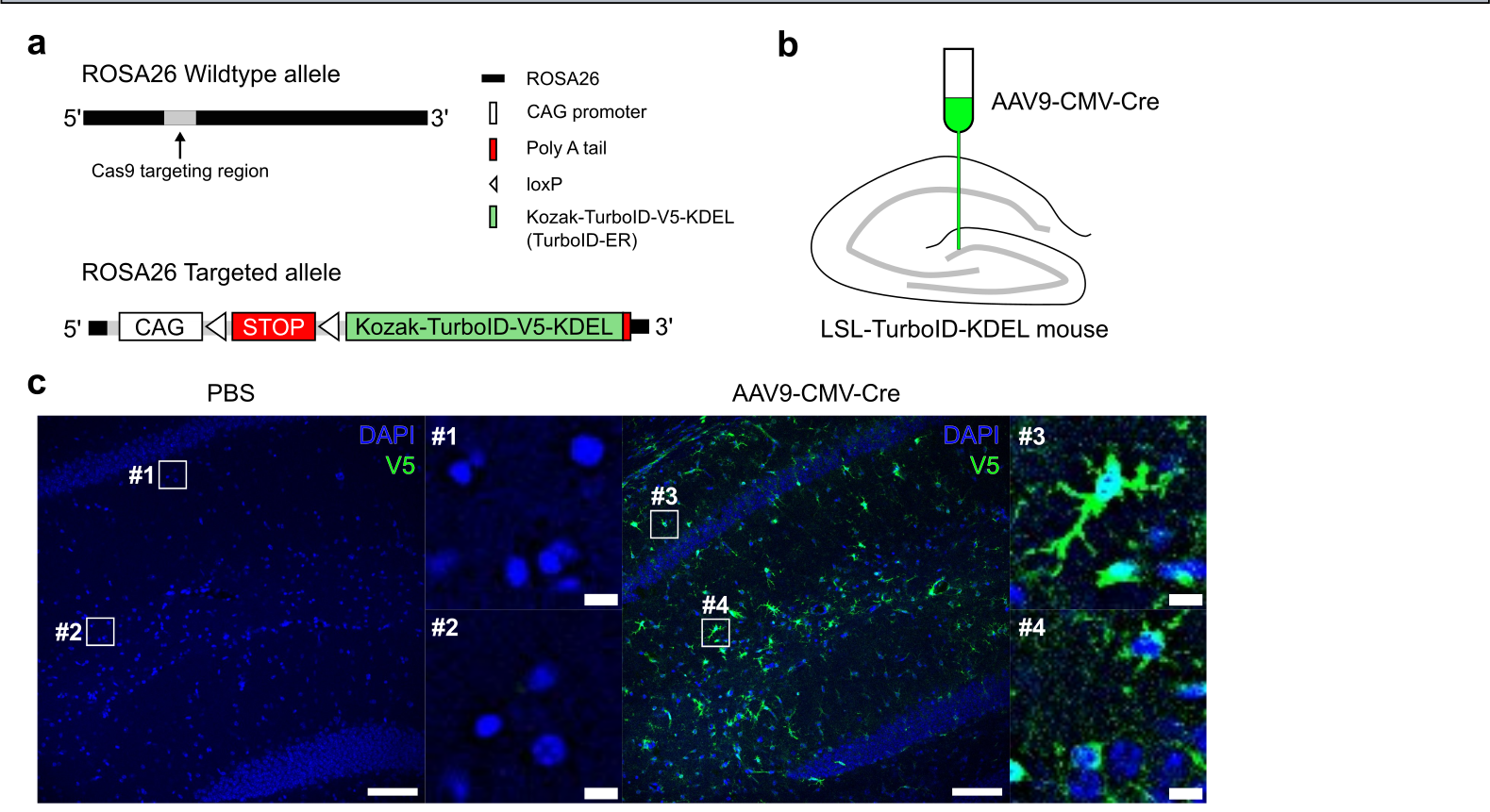

**Extended Data Fig 1. Cre-dependent secretory protein labeling system, LSL-TurboID-KDEL mouse.**

a. Schematic diagram showing the LSL-TurboID-KDEL construct inserted into the ROSA26 locus in C57BL/6 mice. b. Illustration of stereotaxic injection of AAV9-CMV-Cre virus. c. Representative images showing Cre-dependent expression of TurboID-V5. scale bar: 100 μm, insert: 15 μm.

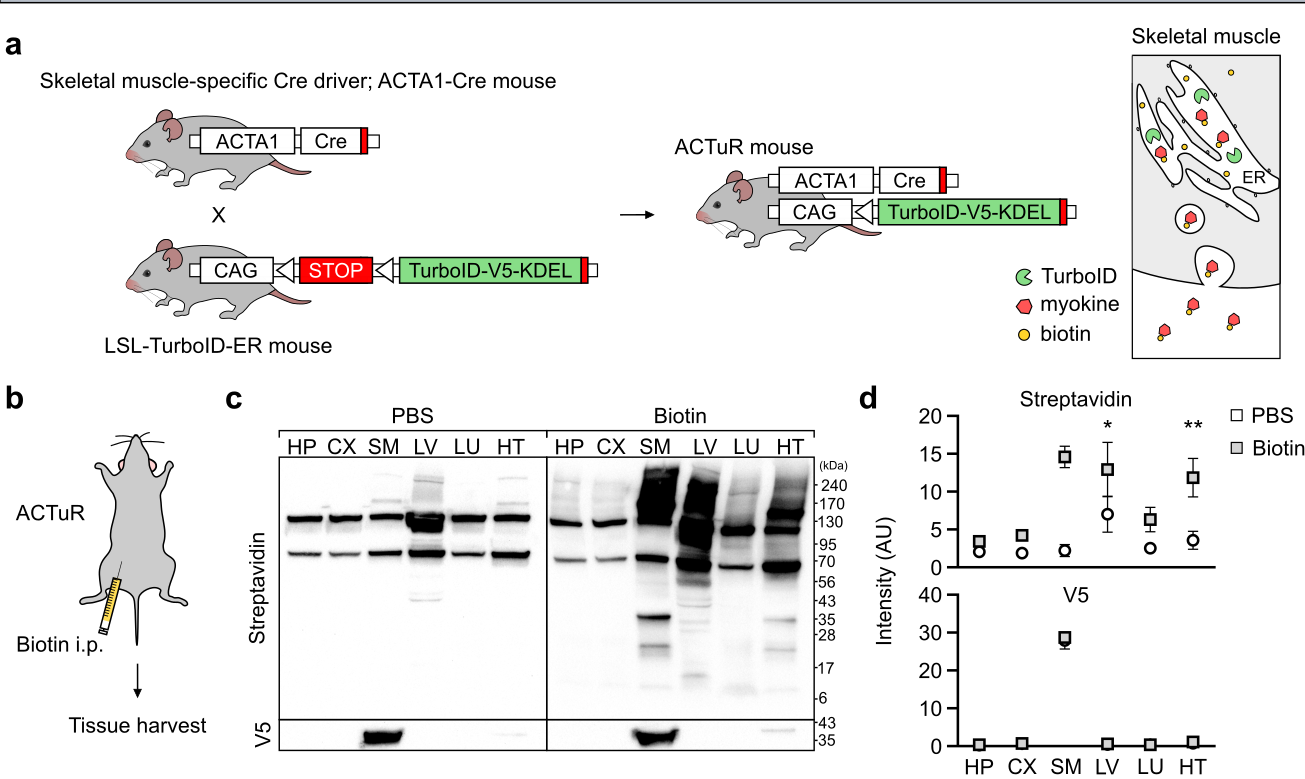

**Extended Data Fig 2. Extended data Figure 2. TurboID-ER is selectively expressed in skeletal muscle of ACTuR mouse.**

a. Schematic for conducting ACTuR mouse model (left). Illustration of secretion of biotinylated myokines from skeletal muscle in ACTuR mouse (right). b. Biotin was intraperitoneally injected for 6 days. c. Representative western blot images showing biotinylated myokines and TurboID-V5 in tissue lysates. d. Bar graph showing the average of intensity of biotinylated proteins (top) and V5 (bottom) of tissue lysates. *p=0.0423, **p=0.0016, unpaired two-tailed Student’s t-test. HP: hippocampus, CX: cortex, SM: skeletal muscle (hind limb), LV: liver, LU: lung, HT: heart. All the data in the bar graphs are presented as the mean ± standard error of the mean (SEM).

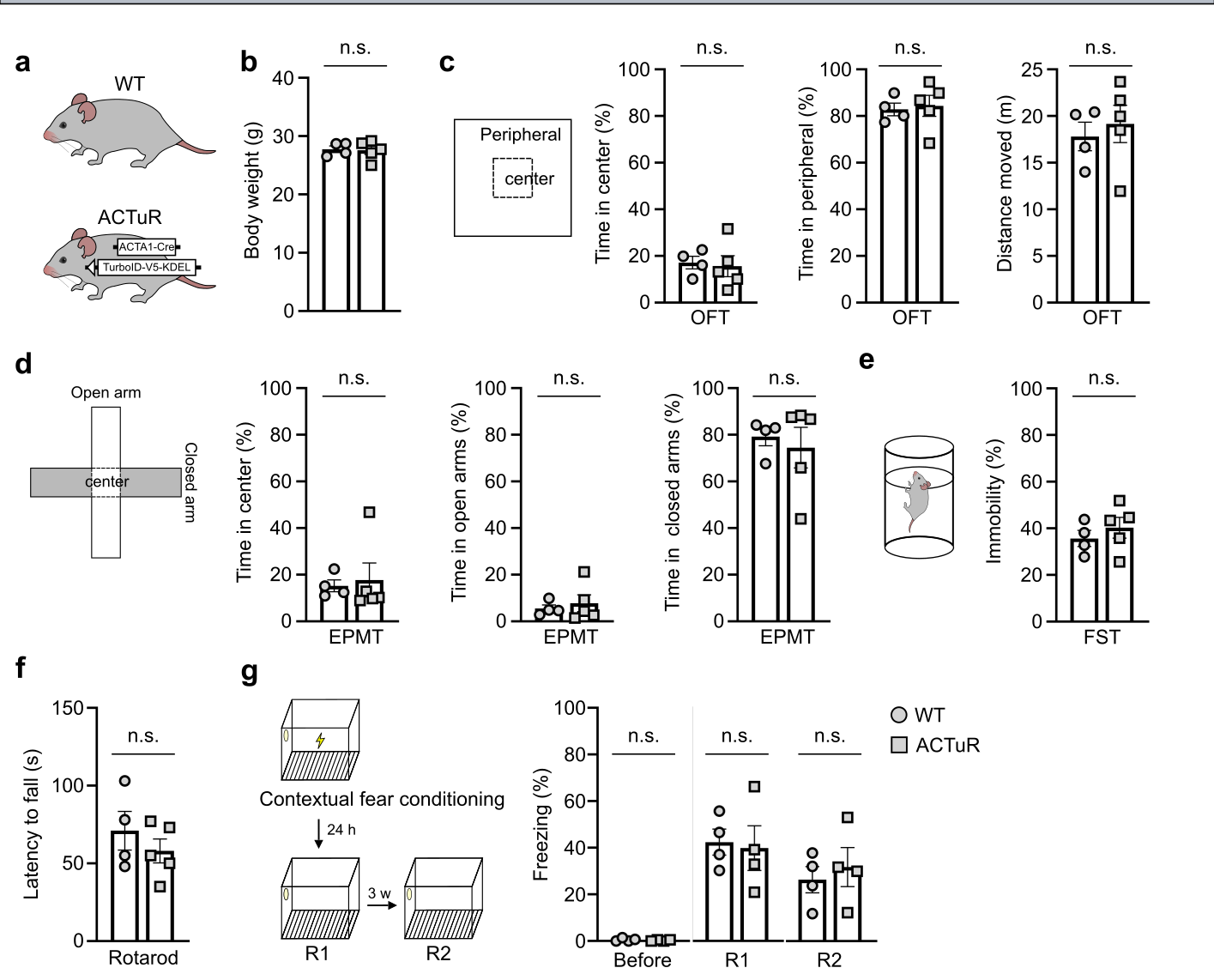

**Extended data Figure 3. No differences between wild-type mouse and ACTuR mouse in diverse behaviors.**

a. Wild-type (WT) mouse and ACTuR mouse. b. body weight (g) c. Time spent in center (%) and peripheral (%) and distance moved (m) in open field test. d. Time spent in center (%) and open arms (%) and closed arms (%) in elevated plus maze test. e. Immobility (%) in forced swimming test. f. Latency to fall (s) in rotarod test. g. freezing time (%) in contextual fear conditioning (CFC) test. p>0.05, unpaired two-tailed Student’s t-test. All the data in the bar graphs are presented as the mean ± SEM. Each dot in the graphs represents a tested mouse. Each dot in the graphs represents a tested mouse.

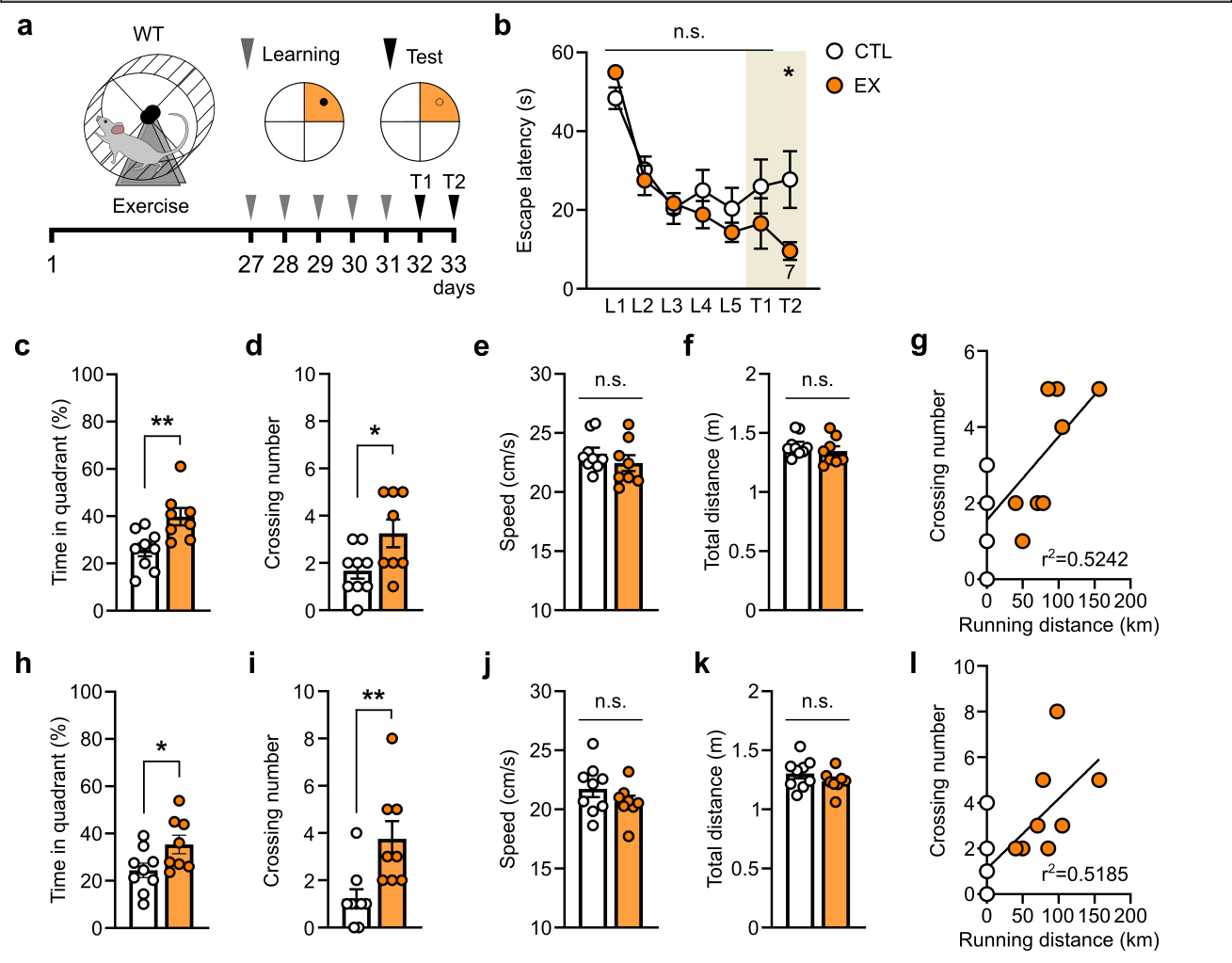

**Extended data Fig 4. Exercise enhances hippocampal memory in Morris water maze test.**

a. Schematic of exercise-induced cognitive analysis. b. Latency to escape (s) during learning and test phase. *p=0.0212, unpaired two-tailed Student’s t-test. CTL: sedentary WT mice; EX: exercised WT mice. c-g. Results in test 1. c. Time spent in quadrant (%). **p=0.0069, unpaired two-tailed Student’s t-test. d. Crossing number. *p=0.0294, unpaired two-tailed Student’s t-test. e. Swimming speed (cm/s). f. Total swimming distance (m). g. Linear regression of crossing number on running distance. h-l. Results in test 2. h. Time spent in quadrant (%). *p=0.0414, unpaired two-tailed Student’s t-test. i. Crossing number. **p=0.0078, unpaired two-tailed Student’s t-test. j. Swimming speed (cm/s). k. Total swimming distance (m). l. Linear regression of crossing number on running distance. All the data in the bar graphs are presented as the mean ± SEM. Each dot in the graphs represents a tested mouse. Each dot in the graphs represents a tested mouse.

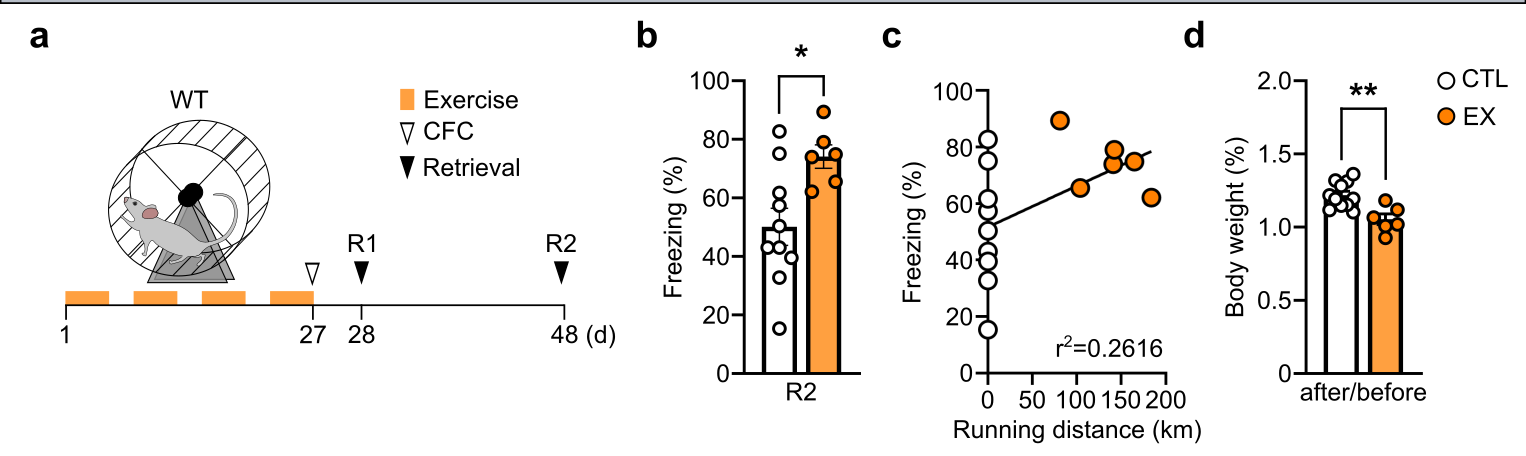

**Extended Data Fig 5. Exercise-induced improvement of hippocampal memory remained for three weeks.**

a. Timeline of CFC test. b. Freezing time (%) three weeks after CFC (R2). *p=0.0163, unpaired two-tailed Student’s t-test. CTL: sedentary WT mice; EX: exercised WT mice. c. Linear regression of freezing time (%) on running distance. d. Ratio of body weight before exercise to body weight after 4 weeks of exercise. **p=0.0023, unpaired two-tailed Student’s t-test. All the data in the bar graphs are presented as the mean ± SEM. Each dot in the graphs represents a tested mouse. Each dot in the graphs represents a tested mouse.

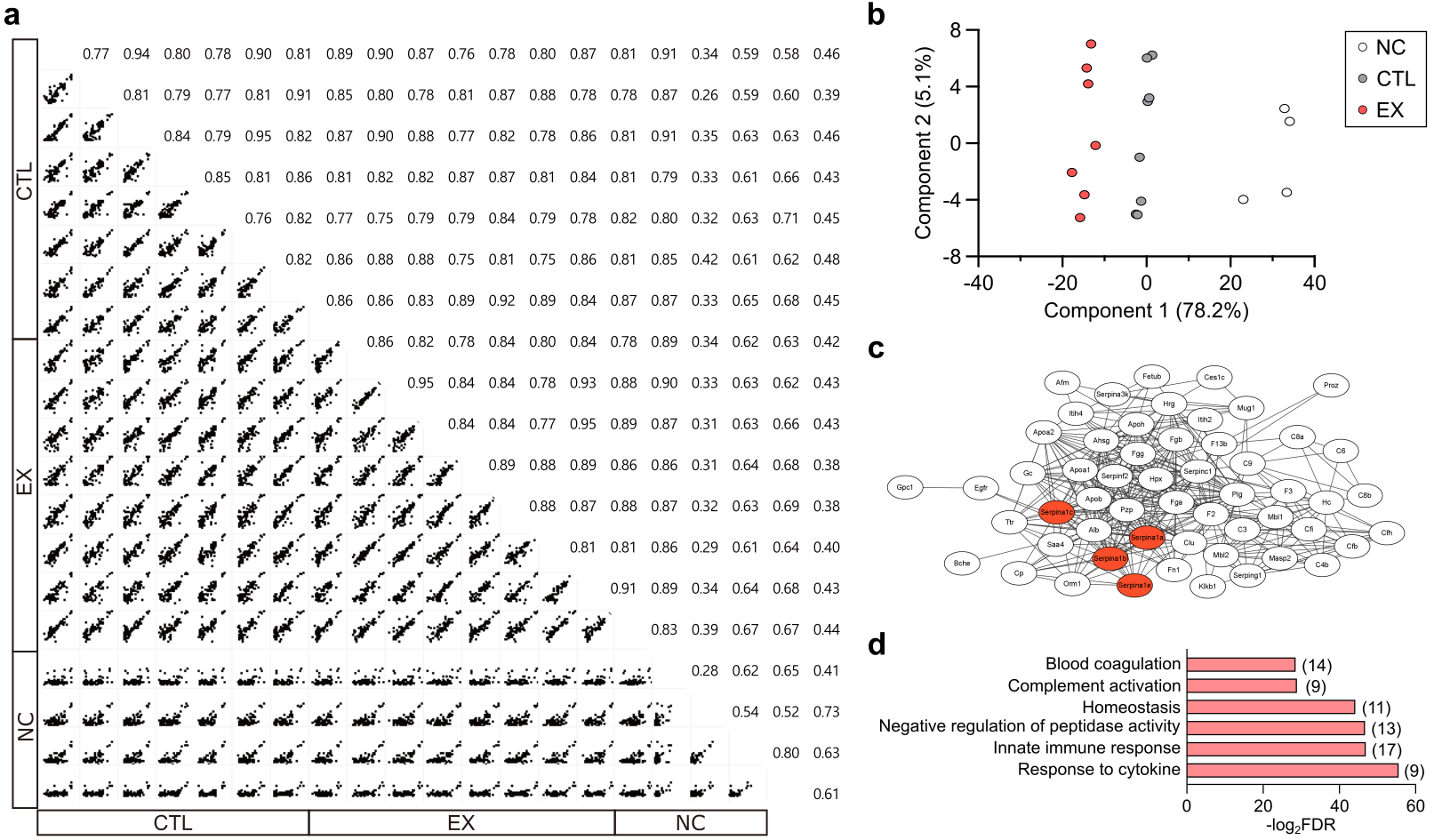

**Extended Data Fig 6. Identification of exercise-induced myokines in plasma.**

a. Pairwise comparison of biotinylated protein quantification and Pearson correlation results. NC: PBS injected sedentary ACTuR mice; CTL: Biotin injected sedentary ACTuR mice; EX: Biotin injected exercised ACTuR mice. b. Principal component analysis of myokines in plasma from each group. c. STRING network analysis of upregulated myokines postexercise. d. GO analysis: biological process (BP) of upregulated myokines postexercise.

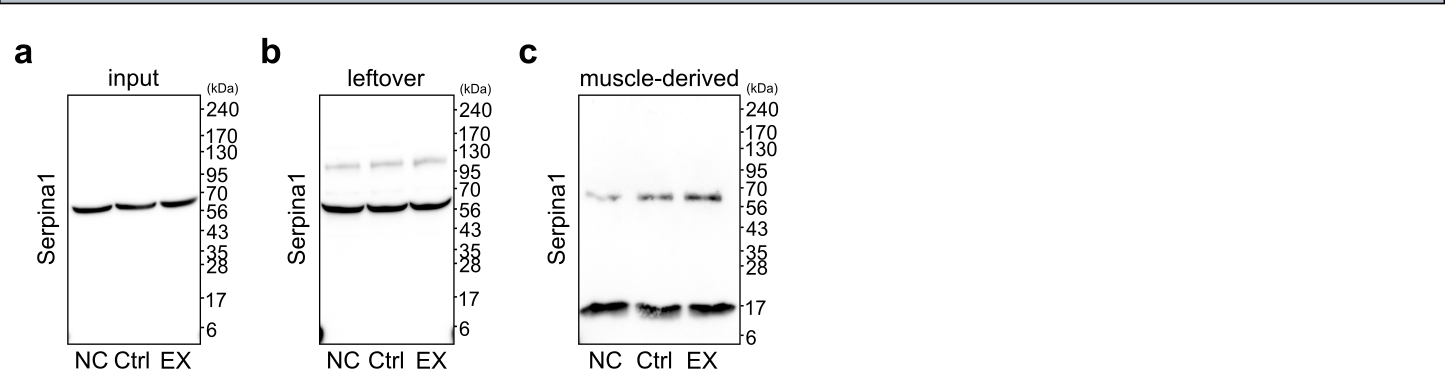

**Extended Data Fig 7. Exercise increases skeletal muscle-derived Serpina1 level in plasma.**

a-c. Representative western blot full images showing Serpina1 level in plasma in input (a), leftover (b), muscle-derived (c) fraction. NC: PBS injected sedentary ACTuR mice; CTL: Biotin injected sedentary ACTuR mice; EX: Biotin injected exercised ACTuR mice.

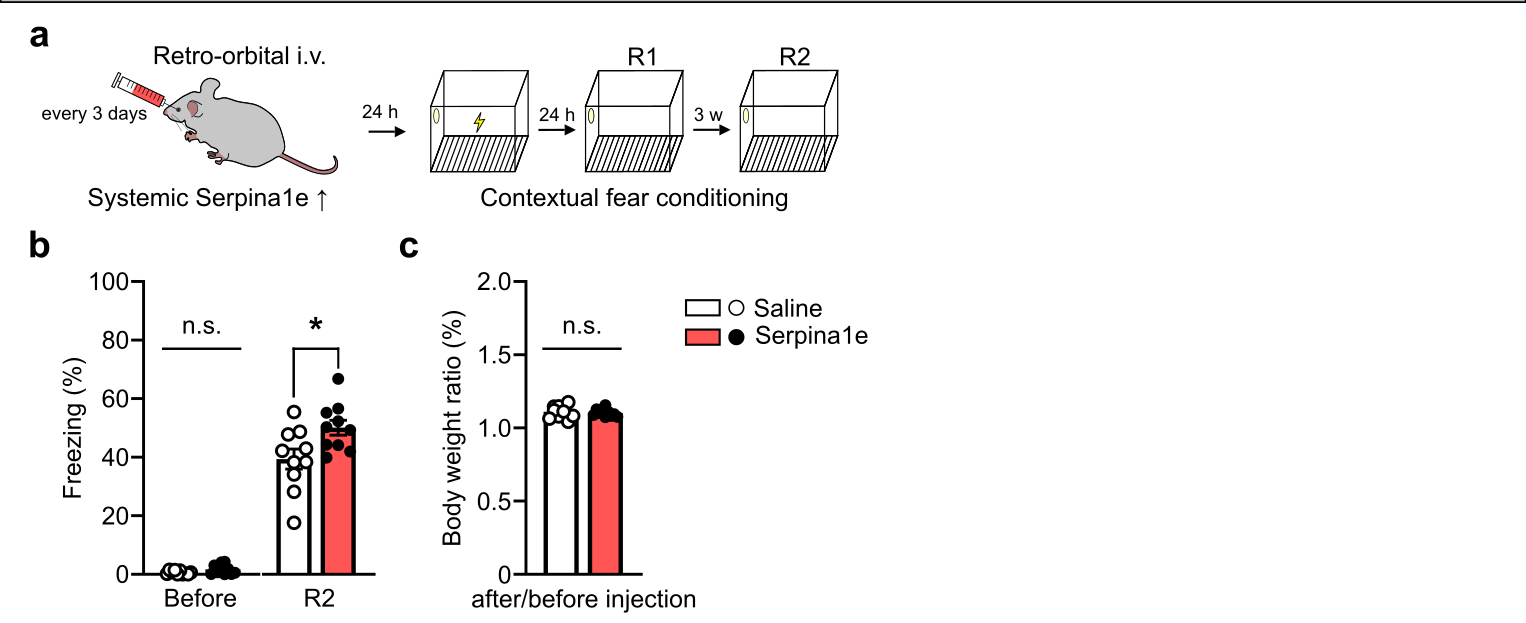

**Extended data Fig 8. Systemic Serpina1e-induced hippocampal memory improvement last for three weeks.**

a. Schematic of systemic administration of recombinant Serpina1e-V5 and CFC test. b. Freezing time (%) in the second retrieval (R2). *p=0.0225, unpaired two-tailed Student’s t-test. Saline: PBS i.v. injected mouse; Serpina1e: Serpina1e i.v. injected mouse. c. Ratio of body weight before intravenous injection to body weight after 4-weeks injection. All the data in the bar graphs are presented as the mean ± SEM. Each dot in the graphs represents a tested mouse. Each dot in the graphs represents a tested mouse.

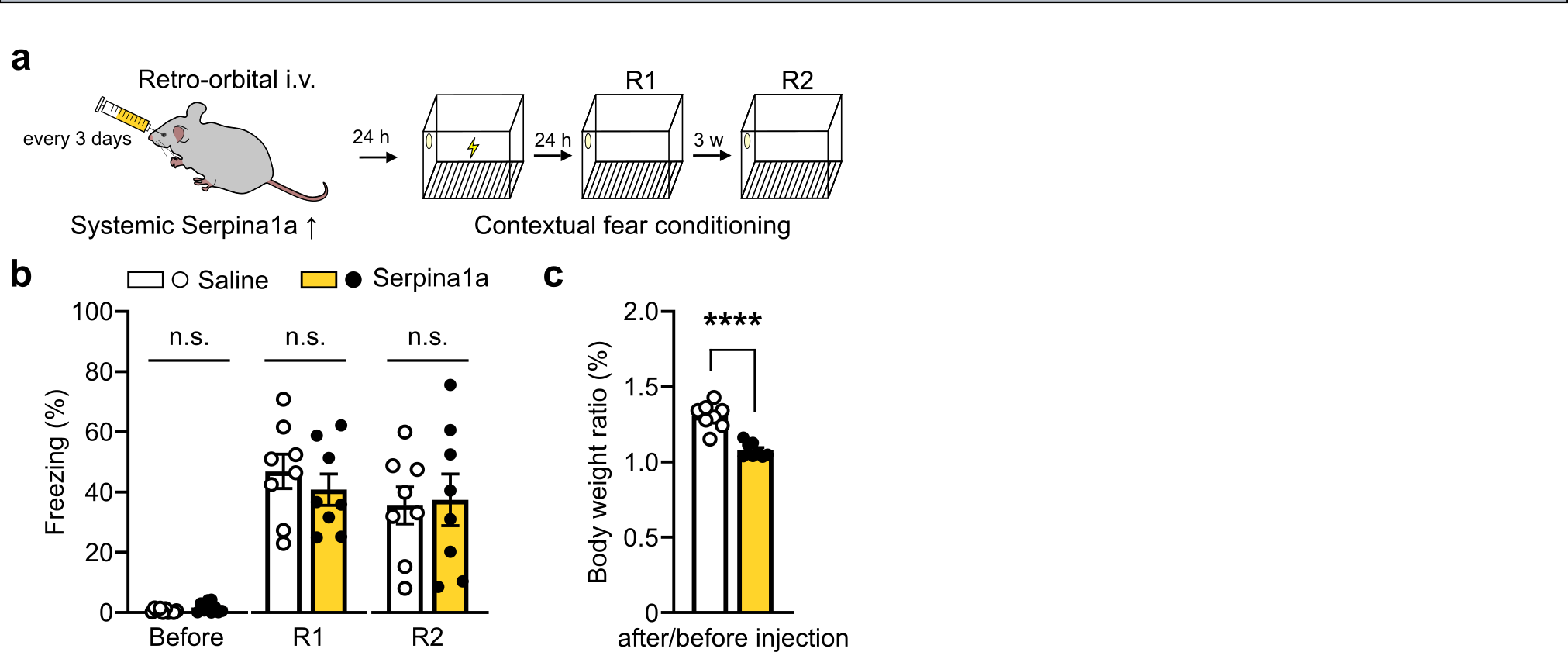

**Extended Data Fig 9. Systemic Serpina1a did not change hippocampal memory.**

a. Schematic of systemic administration of recombinant Serpina1a-V5 and CFC test. b. Freezing time (%) before the CFC and retrieval phase (R1, R2). Saline: PBS i.v. injected mouse; Serpina1e: Serpina1e i.v. injected mouse. c. Ratio of body weight before intravenous injection to body weight after 4-weeks injection. ****p<0.0001, unpaired two-tailed Student’s t-test. All the data in the bar graphs are presented as the mean ± SEM. Each dot in the graphs represents a tested mouse. Each dot in the graphs represents a tested mouse.

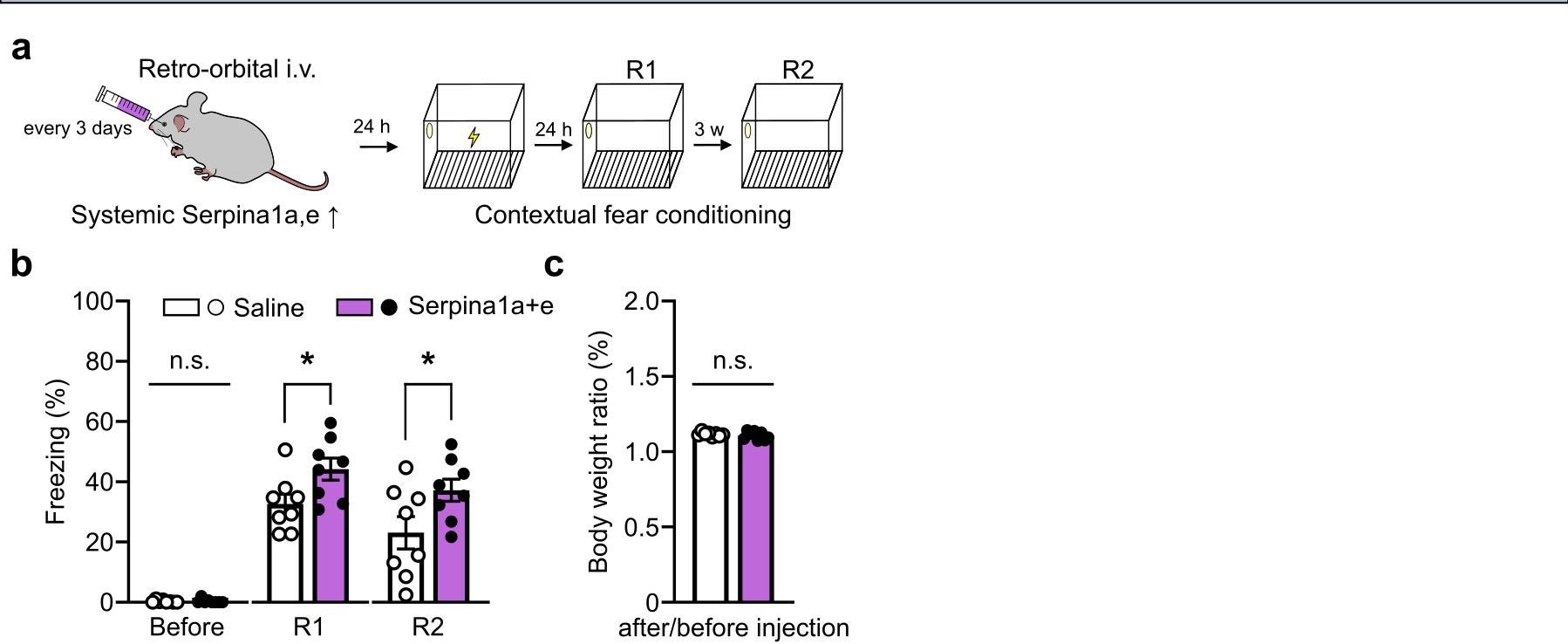

**Extended Data Fig 10. Systemic co-administration of Serpina1a and Serpina1e-induced hippocampal memory improvement that lasts for three weeks.**

a. Schematic of systemic administration of recombinant Serpina1a-V5+Serpina1e-V5 and CFC test. b. Freezing time (%) before the CFC and retrieval phase (R1, R2). *p=0.0324 (R1), *p=0.0474 (R2), unpaired two-tailed Student’s t-test. Saline: PBS i.v. injected mouse; Serpina1a+e: Serpina1a and Serpina1e i.v. injected mouse. c. Ratio of body weight before intravenous injection to body weight after 4-weeks injection. All the data in the bar graphs are presented as the mean ± SEM. Each dot in the graphs represents a tested mouse. Each dot in the graphs represents a tested mouse.

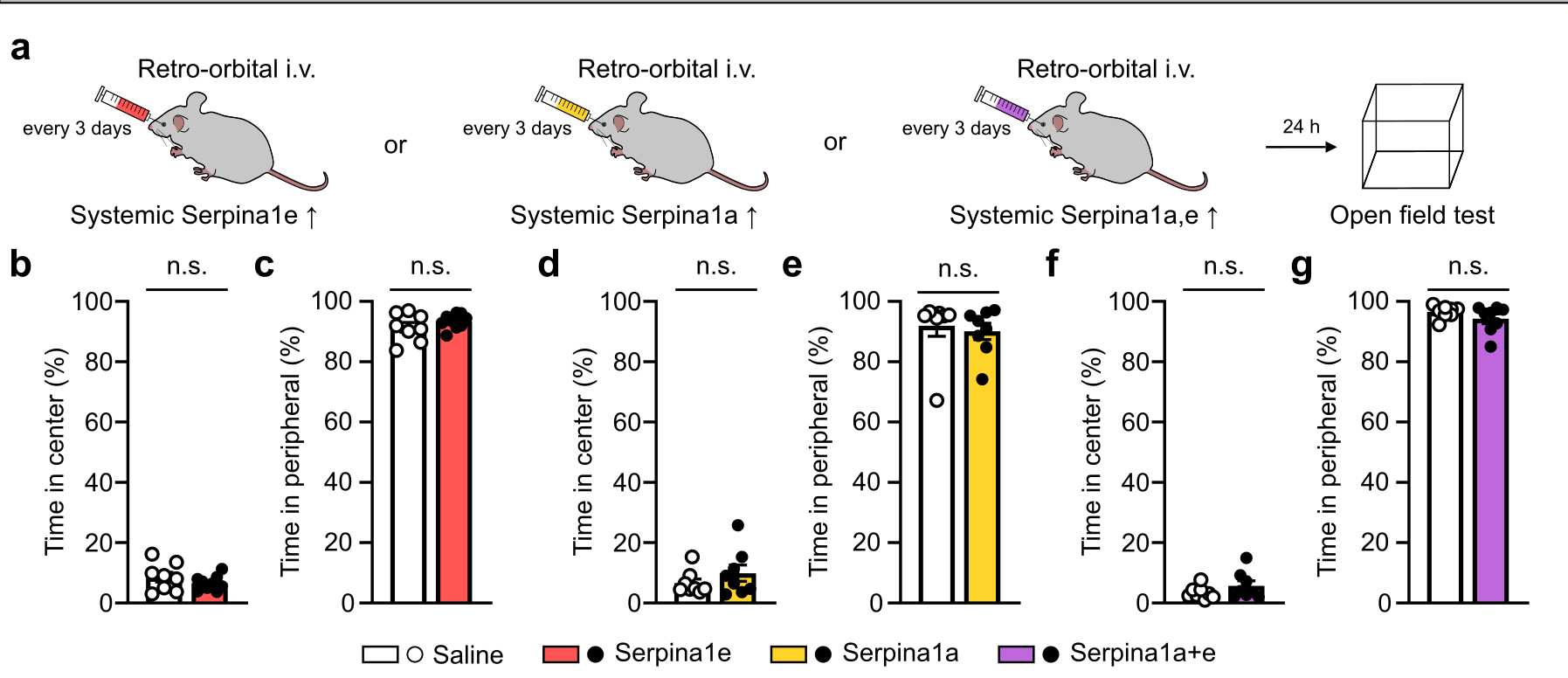

**Extended Data Fig 11. Systemic Serpina1a/Serpina1e did not change general behaviors in open field test.**

a. Schematic of systemic administration of recombinant Serpina1a/Serpina1e and open field test. b-c. Time spent in center (%) and peripheral (%) after intravenous injection for 4-weeks of recombinant Serpina1e. Saline: PBS i.v injected mouse; Serpina1e: Serpina1e i.v. injected mouse; Serpina1a: Serpina1a i.v. injected mouse; Serpina1a+e: Serpina1a and Serpina1e i.v. injected mouse. d-e. Time spent in center (%) and peripheral (%) after intravenous injection for 4-weeks of recombinant Serpina1a. f-g. Time spent in center (%) and peripheral (%) after intravenous injection for 4-weeks of recombinant Serpina1a+Serpina1e. p>0.05, unpaired two-tailed Student’s t-test. All the data in the bar graphs are presented as the mean ± SEM. Each dot in the graphs represents a tested mouse. Each dot in the graphs represents a tested mouse.

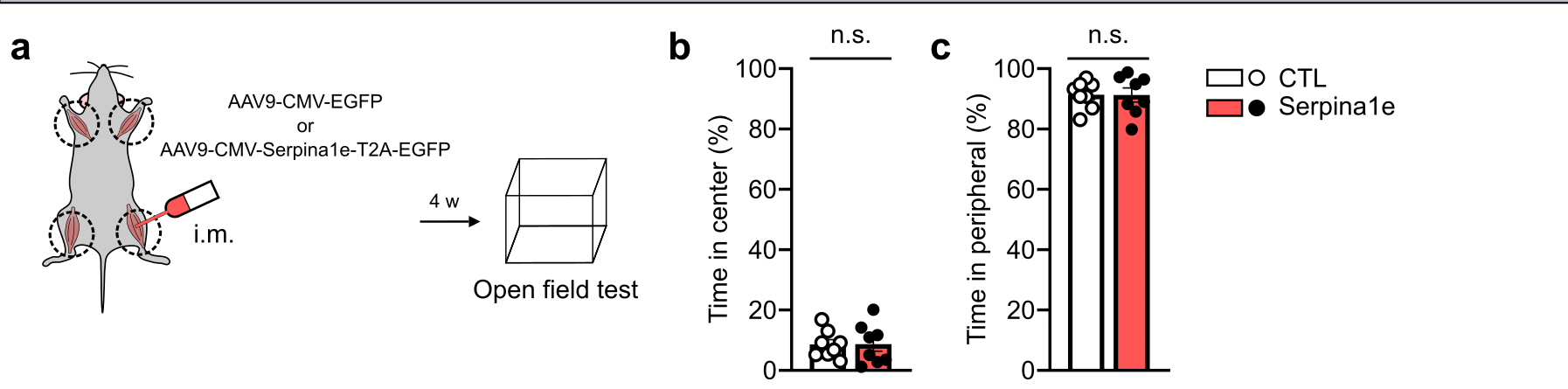

**Extended Data Fig 12. Skeletal muscle-derived Serpina1e improves hippocampal memory improvement in novel object recognition test.**

a. Illustration showing induction of overexpression of Serpina1e in skeletal muscle and open field test. b-c. Time spent in center (%) and peripheral (%) in open field test. CTL: AAV-CMV-EGFP i.m. injected mouse; Serpina1e: AAV9-CMV-Serpina1e-T2A-EGFP i.m. injected mouse. p>0.05, unpaired two-tailed Student’s t-test. All the data in the bar graphs are presented as the mean ± SEM. Each dot in the graphs represents a tested mouse. Each dot in the graphs represents a tested mouse.

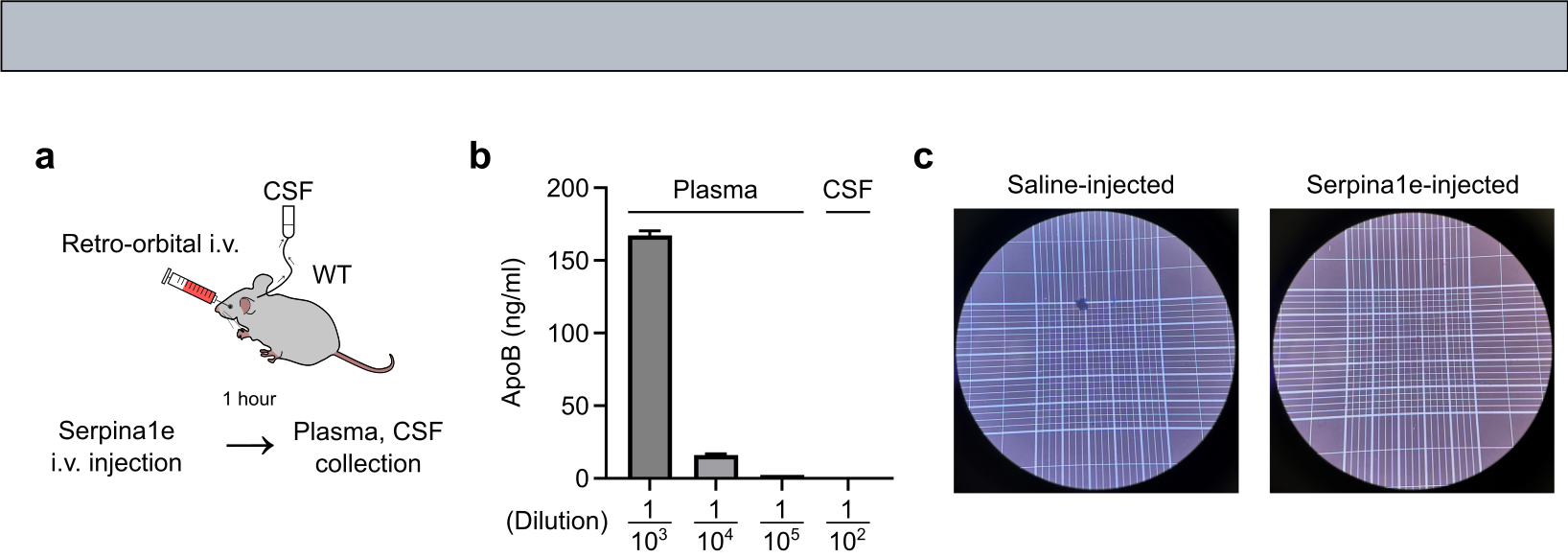

**Extended Data Fig 13. CSF collected from Serpina1e-injected mice showed no signs of blood contamination.**

a. Schematic of collection of CSF and plasma after systemic administration of recombinant Serpina1e. b. ApoB level in plasma and CSF. c. Representative images showing no signs of blood contamination in CSF.

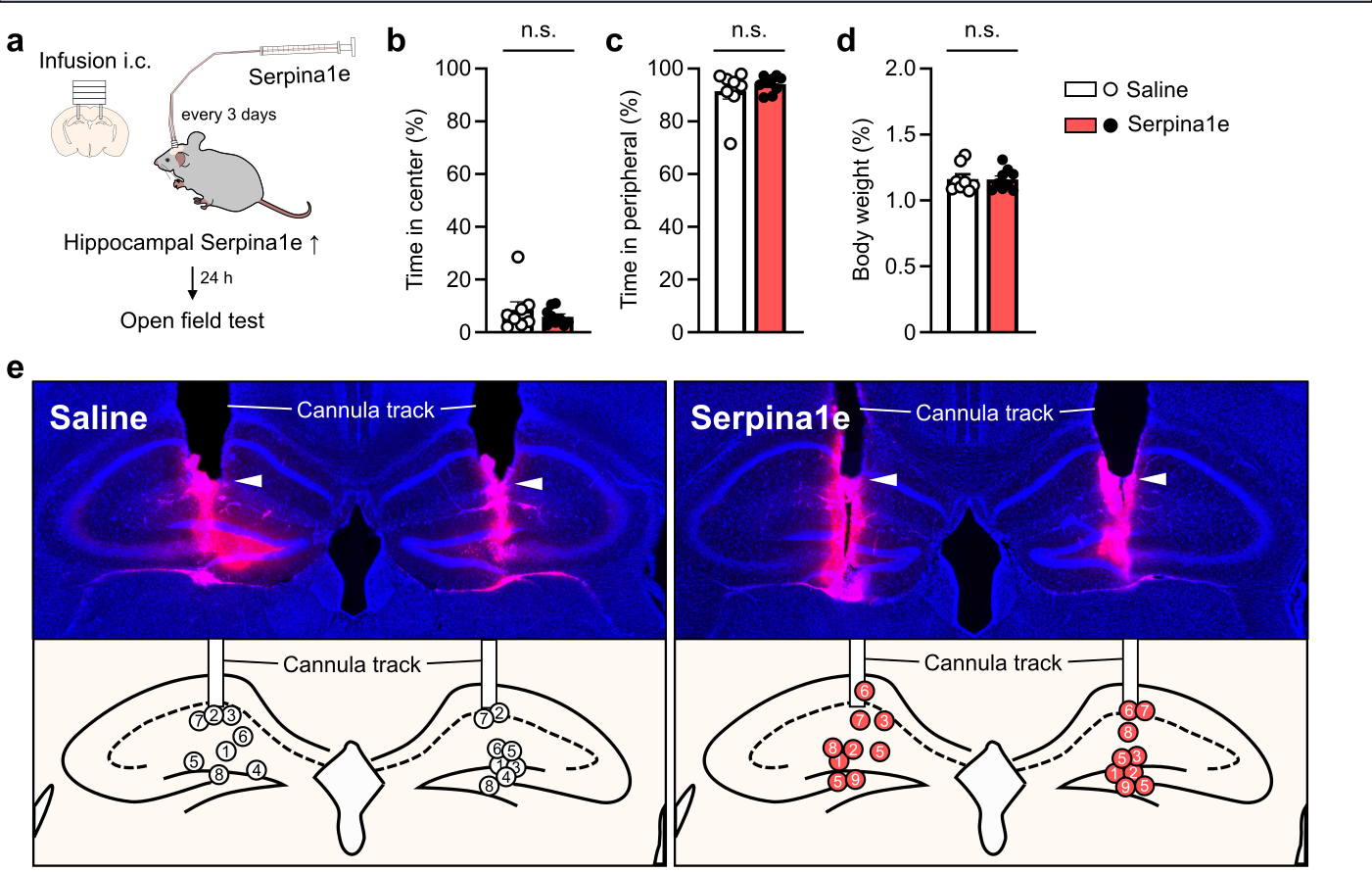

**Extended data Fig 14. Hippocampal infusion of Serpina1e did not change general behaviors in open field test.**

a. Illustration of hippocampal infusion of Serpina1e and behavioral analysis. b-c. Time spent in center (%) and peripheral (%) during open field test. Saline: PBS injected mouse; Serpina1e: and Serpina1e infused mouse. d. Ratio of body weight before infusion into the hippocampus to body weight after 4-weeks hippocampal infusion. p>0.05, unpaired two-tailed Student’s t-test. All the data in the bar graphs are presented as the mean ± SEM. Each dot in the graphs represents a tested mouse. Each dot in the graphs represents a tested mouse. e. Representative images showing cannula tip positions in hippocampus (upper). Illustration of cannula tip location (bottom) and each circle in hippocampus represents each mouse.
