## Extended Data 1 for "Serpina1e mediates the exercise-induced enhancement of hippocampal memory"

| **p-value** | **-log2FC** | protein | | gene |
|---|---|---|---|---|
| 0.722711427 | -0.248962641 | A0A0A6YX | A0A0A6YXQ0_MOUSE | Ighv8-8 |
| 0.403440849 | -0.796122313 | P01872 | IGHM_MOUSE | Ighm |
| 0.993658521 | 0.005143881 | D3YXF5 | D3YXF5_MOUSE | C7 |
| 0.37258521 | 0.304336071 | Q6S9I0 | Q6S9I0_MOUSE | Kng2 |
| 0.75624502 | 0.32279563 | P70663 | SPRL1_MOUSE | Sparcl1 |
| 0.216492472 | 0.439786434 | Q03734 | SPA3M_MOUSE | Serpina3m |
| 0.128142038 | 0.443631649 | Q7TQ48 | SRCA_MOUSE | Srl |
| 0.132701761 | 0.615884304 | E9Q8B5 | E9Q8B5_MOUSE | Cfhr4 |
| 0.451287956 | 0.661767721 | P16301 | LCAT_MOUSE | Lcat |
| 0.46415948 | 0.665898323 | P01820 | HVM44_MOUSE | Hvm44 |
| 0.49724807 | 0.744689703 | Q61702 | ITIH1_MOUSE | Itih1 |
| 0.262984844 | 0.745901346 | Q05920 | PYC_MOUSE | Pc |
| 0.053271605 | 0.784987926 | P97290 | IC1_MOUSE | Serping1 |
| 0.197658166 | 0.813928127 | P01837 | IGKC_MOUSE | Igkc |
| 0.013253846 | 0.879505873 | P01029 | CO4B_MOUSE | C4b |
| 0.043468463 | 0.996493578 | Q9QZF2 | GPC1_MOUSE | Gpc1 |
| 0.043476782 | 1.018636227 | P19221 | THRB_MOUSE | F2 |
| 0.184482813 | 1.040731907 | Q8CG14 | CS1A_MOUSE | C1sa |
| 0.223070038 | 1.067551136 | P06728 | APOA4_MOUSE | Apoa4 |
| 0.071786488 | 1.150080204 | Q60590 | A1AG1_MOUSE | Orm1 |
| 0.02582419 | 1.228719711 | A6X935 | ITIH4_MOUSE | Itih4 |
| 0.024343775 | 1.232150793 | Q61703 | ITIH2_MOUSE | Itih2 |
| 0.084628431 | 1.245183945 | Q06890 | CLUS_MOUSE | Clu |
| 0.025891909 | 1.25657773 | Q91WP6 | SPA3N_MOUSE | Serpina3n |
| 0.275237648 | 1.345273733 | Q9R207 | NBN_MOUSE | Nbn |
| 0.001840601 | 1.346340656 | P09813 | APOA2_MOUSE | Apoa2 |
| 0.001716403 | 1.380518675 | Q61147 | CERU_MOUSE | Cp |
| 0.045261743 | 1.391383648 | O89020 | AFAM_MOUSE | Afm |
| 0.001124145 | 1.397656202 | Q8VCM7 | FIBG_MOUSE | Fgg |
| 0.000344274 | 1.489061356 | Q8K0E8 | FIBB_MOUSE | Fgb |
| 0.080075088 | 1.49221921 | P11276 | FINC_MOUSE | Fn1 |
| 0.192818195 | 1.498591423 | O08677 | KNG1_MOUSE | Kng1 |
| 0.223636028 | 1.502762556 | Q91XL1 | Q91XL1_MOUSE | Lrg1 |
| 4.88E-06 | 1.525296926 | P01027 | CO3_MOUSE | C3 |
| 3.97E-05 | 1.531424999 | Q03311 | CHLE_MOUSE | Bche |
| 5.04E-09 | 1.536142111 | P06909 | CFAH_MOUSE | Cfh |
| 0.000105909 | 1.609607935 | Q01339 | APOH_MOUSE | Apoh |
| 4.84E-06 | 1.642217875 | Q61129 | CFAI_MOUSE | Cfi |
| 0.007137735 | 1.651723623 | Q9QXC1 | FETUB_MOUSE | Fetub |
| 3.27E-06 | 1.667045832 | Q01279 | EGFR_MOUSE | Egfr |
| 0.001071329 | 1.68788147 | P31532 | SAA4_MOUSE | Saa4 |
| 9.85E-07 | 1.693637133 | Q91X72 | HEMO_MOUSE | Hpx |
| 2.62E-05 | 1.695459843 | P04186 | CFAB_MOUSE | Cfb |
| 4.03E-05 | 1.72763896 | P26262 | KLKB1_MOUSE | Klkb1 |
| 3.65E-05 | 1.784654856 | P21614 | VTDB_MOUSE | Gc |
| 1.84E-06 | 1.821790695 | P39039 | MBL1_MOUSE | Mbl1 |
| 1.61E-10 | 1.833254576 | P32261 | ANT3_MOUSE | Serpinc1 |
| 1.01E-05 | 1.848917961 | Q00623 | APOA1_MOUSE | Apoa1 |
| 0.001829124 | 1.865626335 | P41317 | MBL2_MOUSE | Mbl2 |
| 0.018669111 | 1.866205454 | Q07968 | F13B_MOUSE | F13b |
| 0.002894539 | 1.965651751 | E9Q414 | APOB_MOUSE | Apob |
| 0.002123319 | 1.979614735 | Q91WP0 | MASP2_MOUSE | Masp2 |
| 9.88E-05 | 2.00830555 | E9Q6D8 | E9Q6D8_MOUSE | C6 |
| 3.70E-06 | 2.008846521 | P29699 | FETUA_MOUSE | Ahsg |
| 5.44E-11 | 2.010499954 | P07724 | ALBU_MOUSE | Alb |
| 5.91E-06 | 2.040304422 | P06684 | CO5_MOUSE | C5 |
| 2.01E-09 | 2.041594028 | P20918 | PLMN_MOUSE | Plg |
| 0.002139391 | 2.062538624 | P07309 | TTHY_MOUSE | Ttr |
| 0.004134325 | 2.081501961 | Q9DBB9 | CPN2_MOUSE | Cpn2 |
| 1.01E-09 | 2.117223978 | Q61247 | A2AP_MOUSE | Serpinf2 |
| 1.02E-10 | 2.118560791 | P23953 | EST1C_MOUSE | Ces1c |
| 7.05E-10 | 2.165495634 | A0A0R4J0I | A0A0R4J0I1_MOUSE | Serpina3k |
| 0.000223133 | 2.165515184 | Q9CQW3 | PROZ_MOUSE | Proz |
| 1.28E-05 | 2.184031725 | E9PV24 | FIBA_MOUSE | Fga |
| 7.48E-06 | 2.206595898 | P06683 | CO9_MOUSE | C9 |
| 2.77E-10 | 2.266057491 | P42703 | LIFR_MOUSE | Lifr |
| 1.43E-09 | 2.277910709 | Q61838 | PZP_MOUSE | Pzp |
| 2.65E-11 | 2.301855564 | Q921I1 | TRFE_MOUSE | Tf |
| 1.37E-05 | 2.333520889 | P07759 | SPA3K_MOUSE | Serpina3k |
| 8.43E-09 | 2.337373972 | Q8K182 | CO8A_MOUSE | C8a |
| 4.51E-08 | 2.364802122 | P07758 | A1AT1_MOUSE | Serpina1a |
| 3.17E-07 | 2.380060196 | Q8BH35 | CO8B_MOUSE | C8b |
| 8.51E-05 | 2.390520334 | Q9ESB3 | HRG_MOUSE | Hrg |
| 2.90E-09 | 2.440441132 | P28665 | MUG1_MOUSE | Mug1 |
| 2.62E-05 | 2.447719097 | P22599 | A1AT2_MOUSE | Serpina1b |
| 0.000314467 | 2.983365059 | A0A0R4J0) | A0A0R4J0X5_MOUSE | Serpina1c |
| 6.90E-08 | 3.108385324 | Q00898 | A1AT5_MOUSE | Serpina1e |
| 5.23E-08 | 3.655235052 | Q00896 | A1AT3_MOUSE | Serpina1c |
| | not detected | | TurboID | |

| gene | GO:CC term |
|---|---|
| Ighv8-8 | external side of plasma membrane [GO:0009897];immunoglobulin complex, circulating [GO:0042571] |
| Ighm | B cell receptor complex [GO:0019815];cell surface [GO:0009986];cytosol [GO:0005829];external side of plasma membrane [GO:0009897];extracellular space [GO:0005615];immunoglobulin complex, circulating [GO:0042571];integral component of membrane [GO:0016021];membrane [GO:0016020];perinuclear region of cytoplasm [GO:0048471];plasma membrane [GO:0005886] |
| C7 | extracellular region [GO:0005576];membrane attack complex [GO:0005579] |
| Kng2 | collagen-containing extracellular matrix [GO:0062023];extracellular region [GO:0005576];extracellular space [GO:0005615] |
| Sparcl1 | extracellular matrix of synaptic cleft [GO:0098965];extracellular space [GO:0005615];glutamatergic synapse [GO:0098978];synapse [GO:0045202] |
| Serpina3m | extracellular space [GO:0005615] |
| Srl | intracellular membrane-bounded organelle [GO:0043231];sarcoplasmic reticulum lumen [GO:0033018];sarcoplasmic reticulum membrane [GO:0033017] |
| Cfhr4 | extracellular space [GO:0005615] |
| Lcat | extracellular space [GO:0005615];high-density lipoprotein particle [GO:0034364] |
| Hvm44 | external side of plasma membrane [GO:0009897];immunoglobulin complex, circulating [GO:0042571] |
| Itih1 | collagen-containing extracellular matrix [GO:0062023];extracellular matrix [GO:0031012];extracellular region [GO:0005576] |
| Pc | cytoplasm [GO:0005737];mitochondrial inner membrane [GO:0005743];mitochondrial matrix [GO:0005759];mitochondrion [GO:0005739] |
| Serping1 | collagen-containing extracellular matrix [GO:0062023];extracellular space [GO:0005615] |
| Igkc | external side of plasma membrane [GO:0009897];extracellular region [GO:0005576];immunoglobulin complex, circulating [GO:0042571];plasma membrane [GO:0005886] |
| C4b | axon [GO:0030424];dendrite [GO:0030425];extracellular region [GO:0005576];extracellular space [GO:0005615];neuronal cell body [GO:0043025];other organism cell [GO:0044216];synapse [GO:0045202] |
| Gpc1 | anchored component of plasma membrane [GO:0046658];cell surface [GO:0009986];collagen-containing extracellular matrix [GO:0062023];cytosol [GO:0005829];endosome [GO:0005768];extracellular matrix [GO:0031012];extracellular space [GO:0005615];Golgi lumen [GO:0005796];membrane raft [GO:0045121];neuronal cell body [GO:0043025];nucleoplasm [GO:0005654];plasma membrane [GO:0005886];synapse [GO:0045202] |
| F2 | collagen-containing extracellular matrix [GO:0062023];external side of plasma membrane [GO:0009897];extracellular space [GO:0005615] |
| C1sa | extracellular region [GO:0005576];extracellular space [GO:0005615] |
| Apoa4 | cell surface [GO:0009986];chylomicron [GO:0042627];cytosol [GO:0005829];extracellular region [GO:0005576];extracellular space [GO:0005615];high-density lipoprotein particle [GO:0034364];synapse [GO:0045202];very-low-density lipoprotein particle [GO:0034361] |
| Orm1 | extracellular space [GO:0005615] |
| Itih4 | collagen-containing extracellular matrix [GO:0062023];cytoplasm [GO:0005737];extracellular space [GO:0005615];plasma membrane [GO:0005886] |
| Itih2 | collagen-containing extracellular matrix [GO:0062023];extracellular region [GO:0005576] |
| Clu | aggresome [GO:0016235];apical dendrite [GO:0097440];cell periphery [GO:0071944];cell surface [GO:0009986];chromaffin granule [GO:0042583];cytoplasm [GO:0005737];cytoskeleton [GO:0005856];cytosol [GO:0005829];extracellular space [GO:0005615];growth cone [GO:0030426];intracellular membrane-bounded organelle [GO:0043231];mitochondrial inner membrane [GO:0005743];mitochondrion [GO:0005739];neurofibrillary tangle [GO:0097418];neuron projection [GO:0043005];nucleus [GO:0005634];perinuclear endoplasmic reticulum lumen [GO:0099020];perinuclear region of cytoplasm [GO:0048471];protein-containing complex [GO:0032991];spherical high-density lipoprotein particle [GO:0034366];synapse [GO:0045202] |
| Serpina3n | extracellular region [GO:0005576];extracellular space [GO:0005615] |
| Nbn | chromosome, telomeric region [GO:0000781];Mre11 complex [GO:0030870];nuclear inclusion body [GO:0042405];nucleolus [GO:0005730];nucleus [GO:0005634];PML body [GO:0016605];replication fork [GO:0005657];site of double-strand break [GO:0035861] |
| Apoa2 | chylomicron [GO:0042627];cytosol [GO:0005829];extracellular region [GO:0005576];extracellular space [GO:0005615];high-density lipoprotein particle [GO:0034364];spherical high-density lipoprotein particle [GO:0034366];very-low-density lipoprotein particle [GO:0034361] |
| Cp | anchored component of plasma membrane [GO:0046658];extracellular space [GO:0005615];plasma membrane [GO:0005886] |
| Afm | cytoplasm [GO:0005737];extracellular space [GO:0005615] |
| Fgg | blood microparticle [GO:0072562];cell cortex [GO:0005938];cell surface [GO:0009986];collagen-containing extracellular matrix [GO:0062023];cytoplasm [GO:0005737];external side of plasma membrane [GO:0009897];extracellular space [GO:0005615];fibrinogen complex [GO:0005577];platelet alpha granule [GO:0031091];synapse [GO:0045202] |
| Fgb | blood microparticle [GO:0072562];cell cortex [GO:0005938];cell surface [GO:0009986];collagen-containing extracellular matrix [GO:0062023];cytoplasm [GO:0005737];endoplasmic reticulum [GO:0005783];external side of plasma membrane [GO:0009897];extracellular space [GO:0005615];fibrinogen complex [GO:0005577];platelet alpha granule [GO:0031091];synapse [GO:0045202] |
| Fn1 | apical plasma membrane [GO:0016324];basement membrane [GO:0005604];collagen-containing extracellular matrix [GO:0062023];endoplasmic reticulum-Golgi intermediate compartment [GO:0005793];extracellular exosome [GO:0070062];extracellular matrix [GO:0031012];extracellular space [GO:0005615];fibrinogen complex [GO:0005577] |
| Kng1 | collagen-containing extracellular matrix [GO:0062023];extracellular region [GO:0005576];extracellular space [GO:0005615] |
| Lrg1 | extracellular space [GO:0005615];intracellular membrane-bounded organelle [GO:0043231] |
| C3 | cell surface [GO:0009986];extracellular region [GO:0005576];extracellular space [GO:0005615];protein-containing complex [GO:0032991] |
| Bche | endoplasmic reticulum [GO:0005783];endoplasmic reticulum lumen [GO:0005788];extracellular space [GO:0005615];membrane [GO:0016020];nuclear envelope lumen [GO:0005641] |
| Cfh | axon [GO:0030424];cytoplasm [GO:0005737];external side of plasma membrane [GO:0009897];extracellular region [GO:0005576];extracellular space [GO:0005615];mitochondrion [GO:0005739];neuronal cell body [GO:0043025];nucleus [GO:0005634];plasma membrane [GO:0005886] |
| Apoh | cell surface [GO:0009986];chylomicron [GO:0042627];cytoplasm [GO:0005737];extracellular space [GO:0005615];high-density lipoprotein particle [GO:0034364];plasma membrane [GO:0005886];very-low-density lipoprotein particle [GO:0034361] |
| Cfi | extracellular space [GO:0005615];membrane [GO:0016020];nucleus [GO:0005634] |
| Fetub | extracellular region [GO:0005576];extracellular space [GO:0005615] |
| Egfr | apical plasma membrane [GO:0016324];basal plasma membrane [GO:0009925];basolateral plasma membrane [GO:0016323];cell junction [GO:0030054];cell surface [GO:0009986];cytoplasm [GO:0005737];early endosome membrane [GO:0031901];endocytic vesicle [GO:0030139];endoplasmic reticulum membrane [GO:0005789];endosome [GO:0005768];endosome membrane [GO:0010008];Golgi membrane [GO:0000139];integral component of plasma membrane [GO:0005887];membrane [GO:0016020];membrane raft [GO:0045121];multivesicular body, internal vesicle lumen [GO:0097489];nuclear membrane [GO:0031965];nucleus [GO:0005634];perinuclear region of cytoplasm [GO:0048471];plasma membrane [GO:0005886];protein-containing complex [GO:0032991];receptor complex [GO:0043235];synapse [GO:0045202] |
| Saa4 | high-density lipoprotein particle [GO:0034364] |
| Hpx | extracellular space [GO:0005615] |
| Cfb | extracellular space [GO:0005615] |
| Klkb1 | extracellular space [GO:0005615] |
| Gc | axon [GO:0030424];cytoplasm [GO:0005737];cytosol [GO:0005829];extracellular space [GO:0005615];perinuclear region of cytoplasm [GO:0048471] |
| Mbl1 | collagen trimer [GO:0005581];extracellular space [GO:0005615];multivesicular body [GO:0005771];rough endoplasmic reticulum [GO:0005791] |
| Serpinc1 | collagen-containing extracellular matrix [GO:0062023];extracellular space [GO:0005615] |
| Apoa1 | cell surface [GO:0009986];chylomicron [GO:0042627];cytoplasmic vesicle [GO:0031410];cytosol [GO:0005829];discoidal high-density lipoprotein particle [GO:0034365];endocytic vesicle [GO:0030139];extracellular region [GO:0005576];extracellular space [GO:0005615];high-density lipoprotein particle [GO:0034364];intermediate-density lipoprotein particle [GO:0034363];low-density lipoprotein particle [GO:0034362];nucleus [GO:0005634];spherical high-density lipoprotein particle [GO:0034366];very-low-density lipoprotein particle [GO:0034361] |
| Mbl2 | collagen-containing extracellular matrix [GO:0062023];collagen trimer [GO:0005581];extracellular space [GO:0005615];protein-containing complex [GO:0032991] |
| F13b | extracellular region [GO:0005576] |
| Apob | chylomicron [GO:0042627];cytoplasm [GO:0005737];cytosol [GO:0005829];endoplasmic reticulum [GO:0005783];endoplasmic reticulum exit site [GO:0070971];extracellular region [GO:0005576];extracellular space [GO:0005615];high-density lipoprotein particle [GO:0034364];intermediate-density lipoprotein particle [GO:0034363];intracellular membrane-bounded organelle [GO:0043231];lipid droplet [GO:0005811];low-density lipoprotein particle [GO:0034362];mature chylomicron [GO:0034359];neuronal cell body [GO:0043025];very-low-density lipoprotein particle [GO:0034361];vesicle lumen [GO:0031983];vesicle membrane [GO:0012506] |
| Masp2 | extracellular space [GO:0005615] |
| C6 | extracellular space [GO:0005615];membrane attack complex [GO:0005579] |
| Ahsg | collagen-containing extracellular matrix [GO:0062023];extracellular matrix [GO:0031012];extracellular region [GO:0005576];extracellular space [GO:0005615];Golgi apparatus [GO:0005794];protein-containing complex [GO:0032991] |
| Alb | basement membrane [GO:0005604];cytoplasm [GO:0005737];endoplasmic reticulum [GO:0005783];extracellular exosome [GO:0070062];extracellular region [GO:0005576];extracellular space [GO:0005615];Golgi apparatus [GO:0005794];myelin sheath [GO:0043209];protein-containing complex [GO:0032991] |
| C5 | extracellular space [GO:0005615];membrane attack complex [GO:0005579] |
| Plg | cell surface [GO:0009986];collagen-containing extracellular matrix [GO:0062023];extracellular region [GO:0005576];extracellular space [GO:0005615];extrinsic component of external side of plasma membrane [GO:0031232];extrinsic component of plasma membrane [GO:0019897];glutamatergic synapse [GO:0098978];intracellular membrane-bounded organelle [GO:0043231];plasma membrane [GO:0005886];Schaffer collateral - CA1 synapse [GO:0098685] |
| Ttr | extracellular space [GO:0005615];protein-containing complex [GO:0032991] |
| Cpn2 | collagen-containing extracellular matrix [GO:0062023];extracellular matrix [GO:0031012];extracellular space [GO:0005615] |
| Serpinf2 | cell surface [GO:0009986];collagen-containing extracellular matrix [GO:0062023];extracellular space [GO:0005615];fibrinogen complex [GO:0005577] |
| Ces1c | endoplasmic reticulum lumen [GO:0005788];extracellular space [GO:0005615] |
| Serpina3k | extracellular space [GO:0005615] |
| Proz | extracellular region [GO:0005576];extracellular space [GO:0005615] |
| Fga | blood microparticle [GO:0072562];cell cortex [GO:0005938];cell surface [GO:0009986];collagen-containing extracellular matrix [GO:0062023];cytoplasm [GO:0005737];external side of plasma membrane [GO:0009897];extracellular space [GO:0005615];fibrinogen complex [GO:0005577];platelet alpha granule [GO:0031091];rough endoplasmic reticulum [GO:0005791];synapse [GO:0045202] |
| C9 | extracellular space [GO:0005615];membrane attack complex [GO:0005579];other organism cell membrane [GO:0044218];plasma membrane [GO:0005886] |
| Lifr | external side of plasma membrane [GO:0009897];extracellular region [GO:0005576];integral component of membrane [GO:0016021];receptor complex [GO:0043235] |
| Pzp | collagen-containing extracellular matrix [GO:0062023];extracellular region [GO:0005576];extracellular space [GO:0005615] |
| Tf | apical plasma membrane [GO:0016324];basal part of cell [GO:0045178];basal plasma membrane [GO:0009925];basement membrane [GO:0005604];cell surface [GO:0009986];cell tip [GO:0051286];clathrin-coated pit [GO:0005905];cytoplasmic vesicle [GO:0031410];dense body [GO:0097433];early endosome [GO:0005769];endocytic vesicle [GO:0030139];endosome [GO:0005768];extracellular region [GO:0005576];extracellular space [GO:0005615];extrinsic component of external side of plasma membrane [GO:0031232];HFE-transferrin receptor complex [GO:1990712];late endosome [GO:0005770];membrane [GO:0016020];perinuclear region of cytoplasm [GO:0048471];plasma membrane [GO:0005886];recycling endosome [GO:0055037];vesicle [GO:0031982];vesicle coat [GO:0030120] |
| Serpina3k | collagen-containing extracellular matrix [GO:0062023];extracellular space [GO:0005615] |
| C8a | extracellular region [GO:0005576];extracellular space [GO:0005615];membrane attack complex [GO:0005579] |
| Serpina1a | collagen-containing extracellular matrix [GO:0062023];endoplasmic reticulum [GO:0005783];extracellular region [GO:0005576];extracellular space [GO:0005615];Golgi apparatus [GO:0005794];intracellular membrane-bounded organelle [GO:0043231] |
| C8b | extracellular region [GO:0005576];extracellular space [GO:0005615];membrane attack complex [GO:0005579] |
| Hrg | collagen-containing extracellular matrix [GO:0062023];endolysosome [GO:0036019];extracellular region [GO:0005576];phagolysosome membrane [GO:0061474];vesicle [GO:0031982] |
| Mug1 | extracellular space [GO:0005615] |
| Serpina1b | endoplasmic reticulum [GO:0005783];extracellular region [GO:0005576];extracellular space [GO:0005615];Golgi apparatus [GO:0005794];intracellular membrane-bounded organelle [GO:0043231] |
| Serpina1c | extracellular space [GO:0005615] |
| Serpina1e | endoplasmic reticulum [GO:0005783];extracellular space [GO:0005615];Golgi apparatus [GO:0005794];intracellular membrane-bounded organelle [GO:0043231] |
| Serpina1c | endoplasmic reticulum [GO:0005783];extracellular region [GO:0005576];extracellular space [GO:0005615];Golgi apparatus [GO:0005794];intracellular membrane-bounded organelle [GO:0043231] |
